# Generalization and extinction of platform-mediated avoidance in male and female rats

**DOI:** 10.1101/2024.10.17.618811

**Authors:** Alba López-Moraga, Laura Luyten, Tom Beckers

## Abstract

**Background:** Understanding the basis of anxiety-related disorders can be advanced by studying the fear learning mechanisms implicated in the transition from adaptive to maladaptive fear. Individuals with anxiety disorders typically exhibit impaired fear extinction, pervasive avoidance, and overgeneralization of fear. While most research has examined deviations in these fear learning characteristics in isolation, their potential interactions remain understudied. Here, we introduce a modification of the platform-mediated avoidance task and use it to chart avoidance, generalization, and extinction using a single procedure in male and female rats.

**Results:** In the first experiment, we demonstrated that male rats readily acquire avoidance, show a gradient of generalization in a two-day generalization test, and gradually reduce avoidance and fear responding under extinction. In the second experiment, female rats likewise exhibited successful avoidance learning, showed gradual generalization and extinction of defensive behaviors. Across both experiments, interesting sex differences emerged. The third experiment aimed at corroborating these sex differences but showed that they were subtler than expected from the prior separate experiments. Finally, we present an open-source automated system to facilitate the processing of DeepLabCut and SimBA output and obtain reliable results for scoring avoidance and freezing behavior.

**Conclusions:** The modified platform-mediated avoidance task can effectively probe avoidance, generalization, and extinction of fear in male and female rats in a single procedure. Our automated behavioral scoring approach offers researchers an efficient and reproducible method to quantify the defensive behaviors of avoidance, freezing and darting in rats.

## Background

Among psychiatric conditions, anxiety-related disorders are the most common; a systematic review of prevalence studies over 44 countries estimated that anxiety-related disorders had a global prevalence of 7.3% [1,2]. A promising avenue for understanding the underpinnings of anxiety-related disorders lies in identifying fear learning mechanisms implicated in the transition from adaptive to maladaptive fear [3–5]. In particular, individuals with anxiety disorders appear to be characterized by deviations in several fear learning processes, such as extinction, avoidance, and generalization [3,6–8]. Fear conditioning in the laboratory can be used as a translational paradigm to understand the mechanisms at play in the etiology, maintenance, and treatment of anxiety-related disorders, including these three processes [3,5]. Indeed, after fear acquisition, during which a neutral stimulus (Conditioned Stimulus; CS) is paired with an aversive stimulus (Unconditioned Stimulus; US), additional phases can be used to study specific fear learning mechanisms in the laboratory. Avoidance can be investigated by letting specific behavioral responses prevent the occurrence of the US, generalization can be investigated by presenting a generalization stimulus (GS) that resembles the original CS and extinction can be examined by repeatedly presenting the CS without the US [3]. Prior human fear conditioning research has shown that people suffering from anxiety disorders show deficits in extinction learning compared with healthy controls [9,10], exceedingly use avoidance strategies as a coping mechanism [11,12] and exhibit increased generalization after fear learning [13]. To date, however, the roles of impaired extinction, pervasive avoidance, and overgeneralization in the onset or maintenance of anxiety disorders have been predominantly investigated in isolation, with research often focusing on one process at a time. Consequently, our understanding of the interplay and mutual causal influences between these three fear learning processes and their potential role in the transition from adaptive fear to clinical anxiety is limited [3].

In the current study, we address this gap by introducing a new version of the platform-mediated avoidance (PMA) task that allows for assessing avoidance, generalization, and extinction using a single procedure. Although Pavlovian fear conditioning is a translational paradigm that can be applied across species, our research focuses on rats for two important reasons: the possibility (1) to induce a real approach-avoidance conflict and (2) to study sex differences in a highly controlled way.

A real approach-avoidance conflict is established in the PMA procedure as follows. Rats subjected to food restriction are first trained to press a lever in exchange for a food reward. In a subsequent fear acquisition phase, rats are given the opportunity to avoid a CS-signaled foot shock by stepping onto a platform during CS presentation, at the expense of being able to continue lever pressing for food, as the platform is too far from the lever. Thus, food-seeking behavior competes with fearful avoidance [14]. In this task, the cost of avoiding is considerable, which is a translational feature that distinguishes PMA from other active avoidance procedures [15]. This mimics the reality of individuals suffering from anxiety-related disorders who often face approach-avoidance dilemmas and tend to opt for avoidance strategies even when the cost of avoiding is high, a feature that is not well captured by many other lab tasks. Using the PMA task, researchers have recently investigated extinction [14,16–22] and extinction with response prevention [23,24]. However, until now, the procedure has not yet been used to study generalization.

As mentioned above, a second aim of our study was to investigate sex differences, in the absence of human sociocultural factors. The need for biomedical research to counter male bias is pressing. Women are twice as likely to be afflicted as men by anxiety-related disorders [1]. This disparity may have a biological basis that is likely to be evolutionary conserved and easier to dissect in an animal model. Remarkably, there is a reverse bias in preclinical research on anxiety, especially in neuroscience, with many more studies on male subjects than on female subjects only (5.5:1) [25]. In the last decade, funding agencies in Europe and North America have implemented policies to foster the inclusion of both sexes in biomedical research, and prominent journals have published editorials and commentaries providing encouragement and guidance to researchers for designing studies on sex differences and reporting on them in their publications [26–29]. However, an assessment of the influence of these changes in policies over a span of 10 years indicates that while there has been an increase in the inclusion of both sexes across most biological disciplines, studies often still fail to report sample size by sex or perform analyses by sex [30]. Even in clinical research, where formal measures from institutions fostered the increased inclusion of women in clinical trials, studies that include both sexes often fail to provide sex-specific analyses [31]. Ultimately, significant sex differences in etiology and symptomatology, coupled with the lack of attention to these sex differences or even the inclusion of females in research, negatively impact women’s health [32].

Recently, researchers have started investigating differences between male and female rodents in the PMA task. Some authors reported no sex differences in Sprague-Dawley rats [33], whereas others did not report differences in acquisition but only in extinction of PMA in Long-Evans rats [17]. Other authors reported sex differences in acquisition and extinction in C57BL/6J mice [22]. Sex differences in PMA clearly need further scrutiny.

Here, we modified the rodent PMA task to include a two-day generalization phase between the ten-day avoidance acquisition phase and the four-day extinction phase. This allowed us to test avoidance, generalization, and extinction using a single procedure, and investigate if they develop differently in male and female rats. We also introduce a machine-learning approach to automate the analysis of behaviors relevant to the PMA (i.e., avoidance, freezing, and darting).

We hypothesized that rats would readily acquire avoidance across two sessions of avoidance training and that males would show higher freezing than females during this training, in line with previous literature using classical fear conditioning procedures [34–36]. We further hypothesized that there would be a generalization gradient during the second generalization session, and we expected higher generalization in females than in male rats on the basis of prior contextual fear conditioning research [37]. Finally, we hypothesized that rats would gradually extinguish fear to the CS across the fear extinction sessions. Given results observed in other active avoidance tasks [38,39], we predicted that female rats would retain higher avoidance during the extinction sessions than male rats.

## Methods

### Preregistration and data availability

The first experiment reported here, conducted in males only, was not preregistered. For the other two experiments, the design, procedures, sample sizes, and analysis plans were registered on the Open Science Framework (OSF) before the start of data collection. Data and scripts for the three experiments can be found on OSF as well: https://osf.io/ncb8j/.

### Subjects

All experiments were performed in accordance with Belgian and European laws (Belgian Royal Decree of 29/05/2013 and European Directive 2010/63/EU) and the ARRIVE 2.0 [40] guidelines and approved by the KU Leuven animal ethics committee (project license number: 011/2019). The number of animals used was calculated based on previous studies. The first study was conducted in 8-week-old male Sprague-Dawley rats (270-300 g at arrival, Janvier Labs, Le Genest-Saint-Isle, France), the second study was conducted in 8-week-old female Sprague-Dawley rats (200-240 g at arrival, Janvier Labs) and the third study conducted in 8-week-old male (270-300 g at arrival) and female (200-240 g at arrival) rats (Janvier Labs).

Animals were housed in groups of 3 on a 12-hour light-dark cycle (lights on at 7 am), and experiments were performed between 9 am and 4 pm. The cages had bedding and cage enrichment in the form of a red polycarbonate tunnel hanging from the top of the grid. Water was available ad libitum for the entire experiment, except during behavioral testing. Food was provided ad libitum until 1 day before the start of the experiments. From then on, animals were fed ad libitum for an hour after each test session. Animals were habituated to handling for 2 days before the start of the experiment.

### Apparatus

Six identical operant chambers (30.5 cm width, 25.4 cm depth, and 30.5 cm height; Rat Test Cage, Coulbourn Instruments, Pennsylvania, USA) were used simultaneously and were enclosed in sound-attenuating boxes. A 12.8 cm by 15.2 cm platform made of a non-transparent red plastic 3-mm sheet covered approximately ¼ of the grid floor and was always placed in the corner opposite to the lever. Experiments were conducted using Graphic State 3 (Coulbourn Instruments, Pennsylvania, USA) with the house light on. Behavior was continuously recorded during experimental sessions with an IP (Internet Protocol) camera (Foscam C1, Shenzhen, China).

### Lever press training

Lever press training consisted of 1 h of training for 10 consecutive days in Experiments 1 and 2. In Experiment 3, lever press training consisted of 30 minutes of training for 10 consecutive days. Rats were gradually shaped to lever press for grain-based pellets (Experiment 1 and 2: 45 mg 5TUM, TestDiet, St. Louis, MO, USA; Experiment 3: Grain-Based Dustless Precision Pellets® for Rodent, Bio-Serv, Frenchtown, NJ, USA) on a variable interval schedule of reinforcement averaging 30 seconds (VI-30s). Rats that did not reach the criterion of an average of 10 lever presses per minute across the final VI-30 s session were excluded from further analyses (this resulted in the exclusion of three animals in Experiment 3). For Experiment 2, we initially preregistered a criterion of 15 lever presses per minute; however, we lowered this criterion after lever press training to avoid excluding too many rats and in line with the criterion of 10 lever presses per minute in other recent reports using the PMA task [16].

The first training phase consisted of one session of magazine training with the lever retracted, where food pellets were delivered at fixed 2-min intervals. The second phase of lever press training consisted of magazine training with the lever extended, where food pellets were delivered at fixed 2-min intervals, supplemented with direct delivery of a food pellet upon each lever press. This phase lasted one day minimally and required at least 1 lever press/min on average to pass on to the next phase. If there were no lever presses during the first half of the third day of this phase of training, hand shaping was performed. The next phases consisted of lever press training on increasing variable ratio (VR) schedules, progressing from VR 3 through VR 5 and VR 15 to VR 30, where pellets were delivered after a variable number of lever presses, with an increasing average reinforcement criterion per session. On VR 3, a criterion of at least 3 lever press/min was set to pass to the next schedule, a criterion of at least 5 lever press/min for VR 5, a criterion of at least 10 lever press/min for VR 15, and a criterion of at least 15 lever press/min for VR 30. The last phase consisted of training on a VI 30 schedule where the first lever press after a variable refractory period averaging 30 s yielded pellet delivery.

### Experiment 1

Male rats underwent testing in a study that consisted of three phases: avoidance acquisition, generalization testing, and extinction training.

Avoidance acquisition lasted for 10 days in accordance with previous research [14,16,18,41]. On each day, 9 30-s CSs were presented, with an ITI (intertrial interval) averaging 180 s (150 s to 210 s). This ITI was also used in all subsequent sessions. The CS consisted of a pure 3-kHz tone for half of the animals and a 15-kHz tone for the others (counterbalanced). After CS termination, a 2-s, 0.4-mA foot shock US was delivered through the grid floor. If the rats would be on the platform during US presentation, they would not experience the foot shock.

After the 10 days of avoidance acquisition, a two-day generalization test followed. Each generalization session consisted of 3 blocks of 3 trials. Each block started with the original CS and then 2 GSs were presented (GS1: 7 kHz, GS2: 15 kHz, or 3 kHz, depending on CS counterbalancing). The order of the GSs was counterbalanced between animals and test sessions. Each tone lasted for 30 s and no US followed these tone presentations. At the end of each generalization test session, we presented a reminder CS-US pairing to minimize extinction learning. That additional CS is not included in the analyses presented here.

Finally, the experiment concluded with four daily extinction training sessions. Each extinction session consisted of the original 30-s CS being presented nine times, as during the avoidance acquisition session but without US delivery.

### Experiment 2

Female rats were tested in the same procedure as described above. Note that rats were handled by a different researcher for this study, and the two experiments were not conducted in parallel. Other than that, the experimental setup and conditions, the experimental design, and procedures were the same.

### Experiment 3

The experimental design closely mimicked that of Experiments 1 and 2, with a few specific exceptions. First, males and females were tested simultaneously and by the same experimenter. Second, during avoidance training the US co-terminated with the CS rather than occurring at the offset of the CS.

#### Estrous cycle staging

Estrous staging was performed daily for 10 consecutive days after the fourth extinction training session of Experiment 3, to retroactively estimate the estrous stages throughout the behavioral tests without performing the somewhat stressful procedure at the time of these tests. The procedure was performed as described in Marcondes et al., 2002 [42]. In short, rats were brought into a quiet room, different from the experimental test room, where a vaginal smear was collected. Saline (100 µl) was introduced at the entrance of the vaginal canal to flush vaginal cells. The flushed solution was placed on a glass slide and protected with a coverslip for subsequent examination under a light microscope.

### Behavior

Avoidance and freezing during CS presentations were scored manually from the videos. The experimenter was blinded to the test session day and group to the extent possible (shock delivery could give away avoidance training session). Freezing and avoidance were scored as a percentage of CS duration. Avoidance was considered present when at least two paws and the animal’s center of mass were on the platform. Freezing was defined as full immobility except for minimal movements associated with breathing.

A discrimination index of avoidance and freezing during generalization was established by calculating the difference between freezing or avoidance during the presentation of a given tone (i.e., CS) and another tone (i.e., GS1), divided by the sum of freezing or avoidance to both tones [43]. This was done for all the different combinations of CS and GSs.

Lever presses were recorded using Graphic State 3 software (Coulbourn Instruments, Pennsylvania, USA). Lever presses produced 1 min before the tone (pretone) and during the tone were aggregated per block (each block consists of 3 tones), after which the following formula was applied to calculate suppression of lever pressing: (pretone rate – tone rate)/(pretone rate + tone rate) x 100 [44]. Resulting values range from no suppression of lever pressing (0%) to full suppression of lever pressing (100%). For blocks where pretone and tone values were both 0, the rat’s suppression was substituted by the mean group average for that block.

#### DeepLabCut

In addition to the manual scoring of behavior, we also used DeepLabCut (version 2.3.9) for body part tracking in Experiment 3 [45,46]. We labeled 200 frames taken from 10 videos, each video coming from a different animal; 95% of these frames were used for training. Five body parts were selected: nose, left ear, right ear, centroid, and tail base. We used a RestNet-50 neural network [47,48] with default parameters for 300000 training iterations. We validated with 1 shuffle and found that the test error was 3.82 pixels, and the training error was 1.54 pixels (image size: 455 x 256 pixels, downsampled from original 1280 x 720 pixels). This network was used to reanalyze all videos from Experiment 3.

#### SimBA

To further analyze pose estimation data obtained with DeepLabCut, we used SimBA (version 1.96.1) [49]. We created a single animal project configuration with user-defined body parts.

We defined pixels per mm by considering the distance between the width of the bottom of the fear conditioning box.

To analyze avoidance behavior, we defined the platform and adjacent wall sections as the region of interest (ROI). If the centroid body part was inside the ROI, this was considered as avoidance behavior.

We constructed a random forest classifier in SimBA to assess freezing. Four videos were manually annotated; 25876 frames (862.53 s) were annotated with freezing present and 58259 frames (1941.97 s) with freezing absent. The start frame was defined as the first frame in which the animal stopped moving, and the end frame was defined as the last frame before the animal started moving again. We built the model with the following settings: n_estimators = 2000, RF_criterion = gini, RF_max_features = sqrt, RF_min_sample_leaf = 1, under_sample_ratio = 2, class_weights = custom: Freezing present = 2; Freezing Absent = 1, with an 80% training split. The evaluation performance of the freezing classifier had an F1 score of 0.85, precision of 0.83, and recall of 0.87 for freezing present and F1 score of 0.92, precision of 0.93, and recall of 0.91 for freezing absent. We set the discrimination threshold at Pr = 0.65 and a minimum duration of 500 ms.

After that, we performed the cue light analysis add-on from SimBA. In our setup, a dim cue light is on for 2 seconds at the start of any given tone presentation. Using this add-on, we obtain a file with all instances when our light is on. We further processed those data using our own R scripts (see OSF for detailed instructions) to detect when our tone presentations occurred and to obtain freezing and avoidance behavior during CS presentations only.

A subset of videos from SimBA was randomly selected and compared with manual scoring for freezing and avoidance. The intraclass correlation coefficient (ICC) was calculated by comparing the scoring of 9 tone presentations over 10 randomly selected videos between human scorers and the SimBA results. The ICC for the avoidance behavior was 0.996 95% CI [0.994, 0.997] and the ICC for the freezing behavior was 0.941 95% CI [0.912, 0.961]. The videos and all project information of DLC and SimBA used to obtain these values can be found on OSF.

### Data analysis

Statistical analyses of behavior during tone presentation was preregistered for Experiments 2 and 3 (deviations from the preregistration are indicated below). Experiment 1 was not preregistered, but followed the analyses preregistered for Experiment 2. The 9 CSs were grouped in blocks of 3 CSs for acquisition and extinction phases. For the generalization phase a mean of the three tone presentations per session was calculated for the CS as well as for GS1 and GS2. Data were analyzed using (mixed) repeated-measures (RM) ANOVA and significant effects were followed up by post-hoc tests with multiple-test corrections. When assumptions were violated, nonparametric statistical analyses were performed. For the nonparametric mixed repeated measures ANOVAs, we applied an aligned rank transform ANOVA using the ARTool [50] package from R (Version 4.1.3; R Core Team, 2022) [51] and R Studio (Version 9.1.372; RStudio Team, 2021) [52]. All other statistical analyses were performed using afex [53] or rstatix [54]. Data were processed and plotted using the tidyverse [55], reshape [56], and zoo [57] packages.

#### Experiment 1 and 2

To analyze acquisition in Experiments 1 and 2, we used three repeated-measures ANOVAs on avoidance, freezing, and suppression of lever pressing during the first block of sessions 1, 2, and 10. Similar analyses were conducted to evaluate the generalization test, but we included stimulus type (CS, GS1, or GS2) as an additional within-subject measure.

Therefore, generalization was assessed with three two-way (Day, Stimulus) repeated-measures ANOVAs on avoidance, freezing, and suppression of lever pressing. We analyzed extinction learning with three repeated measures ANOVAs comparing avoidance, freezing, and suppression of lever pressing from the first to the fourth session. These analyses were preregistered for Experiment 2, but not for Experiment 1.

As secondary analyses, we preregistered for Experiment 2 the comparison of non-persistent and persistent avoider rats. For Experiment 1 we did the same analysis. A rat was considered a persistent avoider when the time on the platform during the CS was more than 50% during the first block of the first extinction day, a criterion similar to that used in a previous study [16]. We also preregistered that we would consider an animal to be a darter if it would exhibit at least one dart between the third and the seventh CS-US pairing during the first avoidance training session, based on [58]. We additionally preregistered testing sex differences between data to be collected in females (Experiment 2) and data previously collected in males (Experiment 1). We preregistered testing sex differences in avoidance, freezing, and suppression of bar pressing during avoidance training in sessions 1, 2, and 10 with 2 (Sex) x 3 (Session) mixed repeated-measures ANOVAs. We preregistered the same approach to study extinction, comparing sessions 1 to 4. For the generalization test, we added Stimulus type as a within-factor to the above mixed repeated-measures ANOVAs, comparing sessions 1 and 2. As an exploratory analysis, we also calculated a discrimination index for Experiments 1 and 2. Using RM ANOVAs we performed tests to compare the discrimination indices of avoidance and freezing between the three tone comparisons (CS vs GS1, CS vs GS2, and GS1 vs GS2) over sessions 1 and 2.

#### Experiment 3

We preregistered three mixed RM ANOVAs comparing the percentage of avoidance, freezing, and suppression between sexes in the first acquisition block in sessions 1, 2, and 10. We also preregistered a similar analysis for the generalization sessions, adding stimulus type (CS, GS1, or GS2) to the mixed ANOVA and using avoidance, freezing, discrimination index, and suppression of lever pressing as dependent variables. Finally, we preregistered testing sex differences in extinction learning with three mixed ANOVAs to compare avoidance, freezing, and suppression of lever pressing during the first to fourth extinction sessions.

As for Experiment 2, we preregistered the same secondary analyses to compare between non-persistent and persistent avoider rats. Finally, we investigated differences during extinction between distinct estrous phases. We classified the female rats as being into a high vs low estrogen phase and used a t-test to evaluate whether there were differences between the resulting groups in avoidance, freezing, or suppression of lever pressing.

## Results

### A clear gradient of generalization can be obtained in a modified platform-mediated avoidance task in male rats

Experiment 1 introduced a test of generalization between the acquisition and extinction of signaled platform-mediated avoidance in male rats (see Figure 1). Rats were first trained to press a lever for food, after which cued avoidance was acquired. A two-day generalization test was then conducted, after which the experiment concluded with an extinction phase.

**Figure 1.**
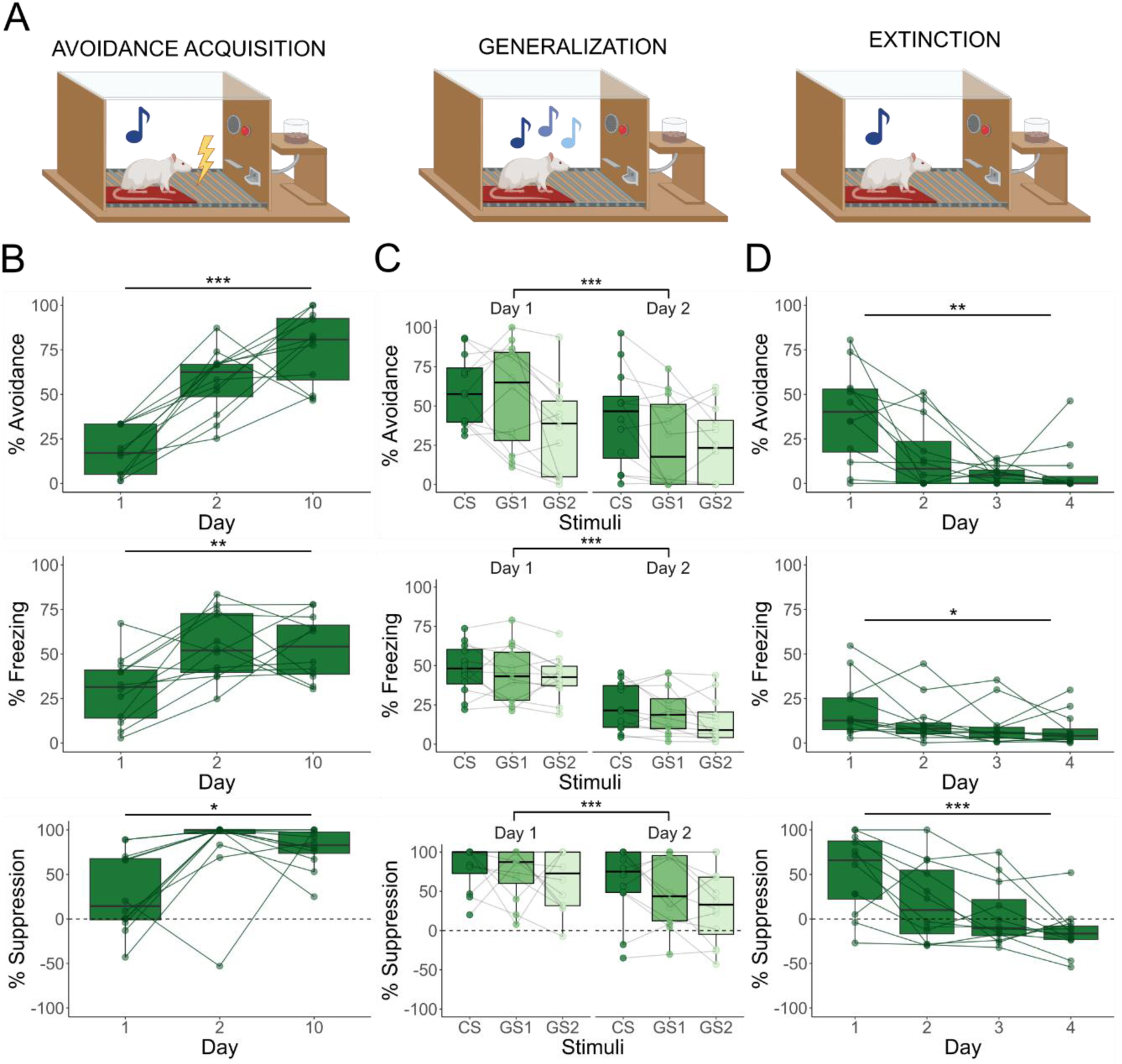
The boxplots represent the average of the first 3 CSs on a given day, except in the generalization test where they represent the average of each stimulus type on a given day. Results are expressed in % of tone duration. **A.** Graphic representation of the avoidance acquisition, generalization, and extinction phases. Different shades of blue denote different tone frequencies in generalization test. **B.** Defensive behavior increased significantly over acquisition sessions: avoidance (F(2, 33) = 35.34, p < 0.001), freezing (F(2, 33) = 6.63, p = 0.004), and suppression of lever pressing (χ^2^ (2) = 9.23, p = 0.01). **C.** During generalization testing, avoidance behavior (F(2, 55) = 8.98, p < 0.001) and lever press suppression (F(2, 55) = 4.75, p = 0.012) but not freezing (F(2, 66) = 1.33, p = 0.272) differed significantly between stimuli. Avoidance was stronger for the CS than for GS2 (t(55) = 4.21, p < 0.001) and for GS1 than GS2 (t(55) = 2.55, p = 0.035). Lever press suppression was stronger during CS than GS2 (t(66) = 2.48, p = 0.041). **D.** Extinction training significantly reduced avoidance (χ^2^ (3) = 15.83, p = 0.001), freezing (χ^2^ (3) = 9.2, p = 0.027), and suppression of lever pressing (F(3, 44) = 6.51, p < 0.001).

An increase in avoidance behavior was observed in male rats over acquisition sessions (F(2, 33) = 35.34, p < 0.001, ω^2^ = 0.66, for further statistics see Table 1), especially between day 1 and 2 (t(33) = –5.594, p < 0.001, d = –2.28) and day 2 and 10 (t(33) = –2.639, p = 0.033, d = – 1.08). Similarly, freezing behavior increased over training sessions (F(2, 33) = 6.63, p = 0.004, ω^2^ = 0.24), both when comparing day 1 and 2 (t(33) = –3.193, p = 0.008, d = –1.30) and day 1 and 10 (t(33) = –3.111, p = 0.01, d = –1.27). Suppression of lever pressing increased over the avoidance acquisition sessions (χ^2^ (2) = 9.23, p = 0.01, W = 0.38), between day 1 and 2 (V = 10, p = 0.021, r = –0.74) and day 1 and 10 (V = 7, p = 0.009, r = – 0.82) as well. To summarize, all fear-related behaviors show similar dynamics over the avoidance acquisition sessions. Rats acquired avoidance behavior, which steadily increased over the acquisition sessions. Their freezing and suppression of lever pressing likewise increased over the first two sessions and then remained stable over subsequent avoidance acquisition days. For more details about the trial-by-trial data, see Supplementary Figure 1a.

**Table 1.** Statistical test results for Experiment 1.

| Test phase | Measured outcome | Statistical Test | Result | Effect size |
| --- | --- | --- | --- | --- |
| Avoidance acquisition | Avoidance | Repeated measures ANOVA | $F(2, 33) = 35.34, p < 0.001$ | $\omega^2 = 0.66 [0.48, 1.00]$ |
| Avoidance acquisition | Avoidance | Pairwise comparison with t-test with Tukey correction | Day 1–2: $t(33) = -5.59, p < 0.001$<br>Day 1–10: $t(33) = -8.23, p < 0.001$<br>Day 2–10 : $t(33) = -2.64, p = 0.033$ | $d = -2.28 [-3.29, -1.27]$<br>$d = -3.36 [-4.54, -2.18]$<br>$d = -1.08 [-1.95, -0.20]$ |
| Avoidance acquisition | Freezing | Repeated measures ANOVA | $F(2, 33) = 6.63, p = 0.004$ | $\omega^2 = 0.24 [0.04, 1.00]$ |
| Avoidance acquisition | Freezing | Pairwise comparison with t-test with Tukey correction | Day 1–2: $t(33) = -3.19, p = 0.008$<br>Day 1–10: $t(33) = -3.11, p = 0.01$<br>Day 2–10 : $t(33) = 0.08, p = 0.996$ | $d = -1.30 [-2.2, -0.41]$<br>$d = -1.27 [-2.16, -0.38]$<br>$d = 0.03 [-0.8, 0.86]$ |
| Avoidance acquisition | Suppression of lever pressing | Friedman test | $\chi^2(2) = 9.23, p = 0.01$ | $W = 0.38 [0.15, 1.00]$ |
| Avoidance acquisition | Suppression of lever pressing | Pairwise comparison with Wilcoxon rank sum test with Bonferroni correction | Day 1–2: $V = 10, p = 0.021$<br>Day 1–10: $V = 7, p = 0.009$<br>Day 2–10 : $V = 43, p = 0.398$ | $r = -0.74 [-0.92, -0.31]$<br>$r = -0.82 [-0.95, -0.48]$<br>$r = 0.3 [-0.32, 0.74]$ |
| Generalization | Avoidance | Non-parametric mixed ANOVA | Stimuli: $F(2, 55) = 8.98, p < 0.001$<br>Day: $F(1, 55) = 23.03, p < 0.001$<br>S*D: $F(2, 55) = 2.68, p = 0.077$ | Stimuli: $\omega_p^2 = 0.06 [0.00, 1.00]$<br>Day: $\omega_p^2 = 0.09 [0.00, 1.00]$<br>S*D: $\omega_p^2 = 0.01 [0.00, 1.00]$ |
| Generalization | Avoidance | Non-parametric pairwise contrasts with Tukey correction | CS–GS1: $t(55) = 1.65, p = 0.233$<br>CS–GS2: $t(55) = 4.21, p < 0.001$<br>GS1–GS2 : $t(55) = 2.55, p = 0.035$ | $d = 0.29 [-0.18, 0.75]$<br>$d = 0.73 [0.25, 1.12]$<br>$d = 0.44 [-0.02, 0.91]$ |
| Generalization | Freezing | Mixed ANOVA | Stimuli: $F(2, 66) = 1.33, p = 0.272$<br>Day: $F(1, 66) = 48.27, p < 0.001$<br>S*D: $F(2, 66) = 0.11, p = 0.899$ | Stimuli: $\omega_p^2 = 0.01 [0.00, 1.00]$<br>Day: $\omega_p^2 = 0.4 [0.25, 1.00]$<br>S*D: $\omega_p^2 = -0.03 [0.00, 1.00]$ |
| Generalization | Suppression of lever pressing | Non-parametric mixed ANOVA | Stimuli: $F(2, 55) = 4.75, p = 0.012$<br>Day: $F(1, 55) = 17.15, p < 0.001$<br>S*D: $F(2, 55) = 1.87, p = 0.163$ | Stimuli: $\omega_p^2 = 0.03 [0.00, 1.00]$<br>Day: $\omega_p^2 = 0.1 [0.02, 1.00]$<br>S*D: $\omega_p^2 = -0.02 [0.00, 1.00]$ |
| Generalization | Suppression of lever pressing | Non-parametric pairwise contrasts with Tukey correction | CS–GS1: $t(66) = 0.76, p = 0.725$<br>CS–GS2: $t(66) = 2.48, p = 0.041$<br>GS1–GS2 : $t(66) = 1.72, p = 0.207$ | $d = 0.22 [-0.24, 0.68]$<br>$d = 0.72 [0.24, 1.19]$<br>$d = 0.49 [0.02, 0.96]$ |
| Extinction | Avoidance | Friedman test | $\chi^2(3) = 15.83, p = 0.001$ | $W = 0.44 [0.27, 1.00]$ |
| Extinction | Avoidance | Pairwise comparison with Wilcoxon rank sum test with Bonferroni correction | Day 1–2: $V = 62$, $p = 0.113$<br>Day 2–3: $V = 43$, $p = 0.126$<br>Day 3–4: $V = 26$, $p = 0.722$<br>Day 1–3: $V = 66$, **$p = 0.004$**<br>Day 1–4: $V = 73$, **$p = 0.005$**<br>Day 2–4: $V = 44$, $p = 0.103$ | $r = 0.88$ [0.61, 0.97]<br>$r = 0.56$ [-0.19, 0.9]<br>$r = 0.16$ [-0.52, 0.71]<br>$r = 1$ [1, 1]<br>$r = 0.87$ [0.59, 0.96]<br>$r = 0.6$ [-0.13, 0.91] |
| Extinction | Freezing | Friedman test | $\chi^2(3) = 9.2$, **$p = 0.027$** | $W = 0.26$ [0.1, 1.00] |
| Extinction | Freezing | Pairwise comparison with Wilcoxon rank sum test with Bonferroni correction | Day 1–2: $V = 58$, $p = 0.147$<br>Day 2–3: $V = 51$, $p = 0.38$<br>Day 3–4: $V = 46$, $p = 0.622$<br>Day 1–3: $V = 69$, **$p = 0.016$**<br>Day 1–4: $V = 74$, **$p = 0.003$**<br>Day 2–4: $V = 58$, $p = 0.151$ | $r = 0.49$ [-0.11, 0.83]<br>$r = 0.31$ [-0.31, 0.74]<br>$r = 0.18$ [-0.43, 0.68]<br>$r = 0.77$ [0.36, 0.93]<br>$r = 0.9$ [0.67, 0.97]<br>$r = 0.49$ [-0.11, 0.83] |
| Extinction | Suppression of lever pressing | Repeated measures ANOVA | $F(3, 44) = 6.51$, **$p < 0.001$** | $\omega^2 = 0.26$ [0.06, 1.00] |
| Extinction | Suppression of lever pressing | Pairwise comparison with t-test with Tukey correction | Day 1–2: $t(44) = 2.34$, $p = 0.105$<br>Day 1–3: $t(44) = 3.08$, **$p = 0.018$**<br>Day 1–4: $t(44) = 4.28$, **$p < 0.001$**<br>Day 2–3: $t(44) = 0.74$, $p = 0.879$<br>Day 2–4: $t(44) = 1.94$, $p = 0.226$<br>Day 3–4: $t(44) = 1.2$, $p = 0.631$ | $d = 0.96$ [0.107, 1.8]<br>$d = 1.26$ [0.39, 2.12]<br>$d = 1.75$ [0.84, 2.65]<br>$d = 0.3$ [-0.52, 1.13]<br>$d = 0.79$ [-0.05, 1.63]<br>$d = 0.49$ [-0.34, 1.32] |

Bold values indicate statistical significance ( $p < 0.05$ ).

During the two-day generalization test, there was a decrease in avoidance behavior over days (F(1, 55) = 23.03, p < 0.001, ω*_p_*^2^ = 0.09) and a significant difference in avoidance per stimulus (F(2, 55) = 8.98, p < 0.001, ω*_p_*^2^ = 0.06). More concretely, rats avoided more during CS than during GS2 presentations (t(55) = 4.21, p < 0.001, d = 0.73) and during GS1 than GS2 (t(55) = 2.55, p = 0.035, d = 0.44), whereas no differences were found between CS and GS1 (t(55) = 1.65, p = 0.233, d = 0.29). We observed a similar pattern for suppression of lever pressing, with less suppression on the second than on the first day (F(1, 55) = 17.15, p < 0.001, ω _p_^2^ = 0.1) and significant differences in suppression between stimuli (F(2, 55) = 4.75, p = 0.012, ω _p_^2^ = 0.03). Again, there was significantly more suppression during CS than during GS2 presentations (t(66) = 2.48, p = 0.041, d = 0.72), and no difference between CS and GS1 presentations (t(66) = 0.76, p = 0.725, d = 0.22) or GS1 and GS2 presentations (t(66) = 1.72, p = 0.207). Freezing behavior showed a slightly different picture. We observed a significant decrease in freezing from day 1 to day 2 (F(1, 66) = 48.27, p < 0.001, ω*_p_*^2^ = 0.4), but no significant differential freezing depending on stimulus type (see additional statistics in Table 1). We also calculated discrimination indices, which can be found in Supplementary Figure 2 and Supplementary Table 1. Overall, rats showed an orderly generalization gradient in suppression of lever pressing and avoidance. We observed clearly different responding between the CS and the more perceptually dissimilar GS2, and intermediate responding to GS1. In contrast, the generalization curve was flatter for freezing, showing similar responding to the three stimuli. We conclude that the introduction of a generalization phase between avoidance and extinction of platform-mediated avoidance was successful.

After the generalization phase, four extinction sessions followed. Rats successfully extinguished avoidance (χ^2^ (3) = 15.83, p = 0.001, W = 0.44), freezing (χ^2^ (3) = 9.2, p = 0.027, W = 0.26) and suppression of lever pressing (F(3, 44) = 6.51, p < 0.001, ω^2^ = 0.26) over four days of extinction. In fact, all behavioral indices were already significantly reduced by the third extinction session (avoidance: V = 66, p = 0.004, r = 1; freezing: V = 69, p = 0.016, r = 0.77; suppression of lever pressing: t(44) = 3.08, p = 0.018, d = 1.26).

### Assessing the modified platform-mediated avoidance task in female rats

Experiment 2 assessed whether the adapted platform-mediated avoidance task that we introduced in Experiment 1 would yield comparable results in female rats (Figure 2).

**Figure 2.**
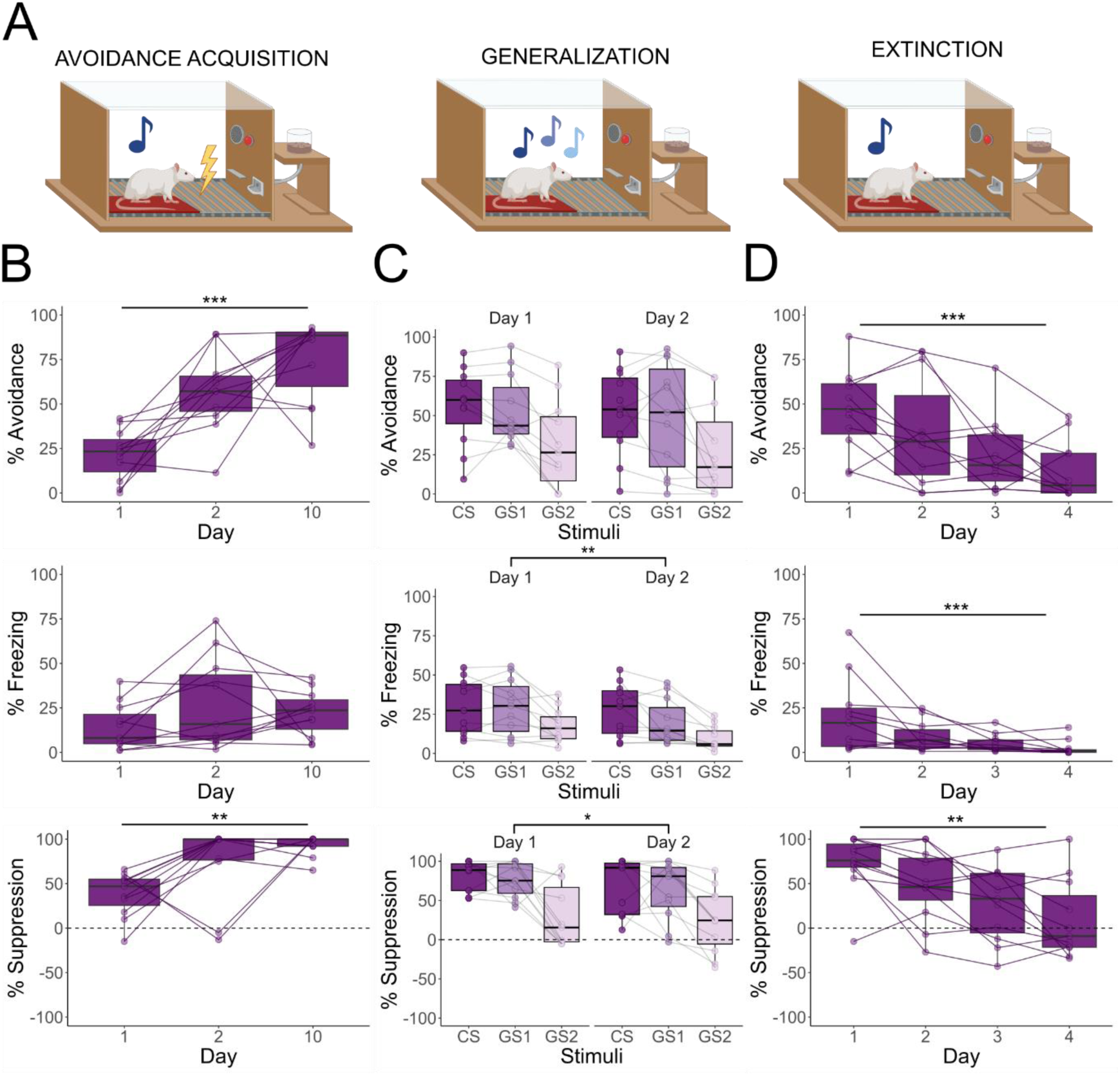
The boxplots represent the average of the first 3 CSs on a given day, except in the generalization test where they represent the average of each stimulus type on a given day. Results are expressed in % of tone duration. **A.** Graphic representation of the avoidance acquisition, generalization, and extinction phases. Different shades of blue denote different tone frequencies in the generalization test. **B.** During avoidance acquisition, an increase in avoidance (F(2, 30) = 19.64, p < 0.001) and suppression of lever pressing (χ^2^ (2) = 11.49, p = 0.003) is observed over the avoidance acquisition sessions, but not in freezing (F(2, 30) = 1.60, p = 0.218). **C.** Defensive behaviors differed significantly between stimuli over the generalization sessions: avoidance (F(2, 60) = 5.2, p = 0.008), freezing (F(2, 50) = 13.02, p < 0.001) and suppression of lever pressing (F(2, 60) = 11.84, p < 0.001). Avoidance (t(60) = 2.93, p = 0.013), freezing (t(50) = 4.84, p < 0.001) and suppression of lever pressing (t(60) = 4.39, p < 0.001) were stronger for CS than GS2. Defensive behaviors also differed between GSs; avoidance (t(60) = 2.63, p = 0.028), freezing (t(50) = 3.82, p = 0.001) and suppression of lever pressing (t(60) = 4.01, p < 0.001) were stronger for GS1 than GS2. **D.** Extinction training reduced avoidance behavior (χ^2^ (3) = 17.61, p < 0.001), freezing (χ^2^ (3) = 21.27, p < 0.001), and suppression of lever pressing (F(3, 40) = 5.32, p = 0.003).

Female rats readily learned to step onto the platform to avoid shock over the 10 days of acquisition (F(2, 30) = 19.64, p < 0.001, ω^2^ = 0.53, see further statistics in Table 2). They avoided significantly more on day 2 than day 1 (t(30) = –4.105, p = 0.008, d = –1.75) and on day 10 than day 1 (t(30) = –6.155, p < 0.001, d = –2.62), with no significant increase from day 2 to day 10. In contrast to the male rats of Experiment 1, female rats did not show a significant increase in freezing behavior over the acquisition sessions. They did show a significant increase in suppression of lever pressing over acquisition sessions (χ^2^ (2) = 11.49, p = 0.003, W = 0.52). This increase was not significant from the first to the second session of acquisition, but it was from the first to the tenth session (V = 0, p < 0.001, r = –1) and from the second to the tenth session (V = 6, p = 0.032, r = –0.78).

**Table 2.** Statistical test results for Experiment 2.

| Test phase | Measured outcome | Statistical Test | Result | Effect size |
| --- | --- | --- | --- | --- |
| Avoidance acquisition | Avoidance | Repeated measures ANOVA | $F(2, 30) = 19.64, p < 0.001$ | $\omega^2 = 0.53 [0.3, 1.00]$ |
| Avoidance acquisition | Avoidance | Pairwise comparison with t-test with Tukey correction | Day 1–2: $t(30) = -4.10, p = 0.008$<br>Day 1–10: $t(30) = -6.16, p < 0.001$<br>Day 2–10 : $t(30) = -2.05, p = 0.118$ | $d = -1.75 [-2.74, -0.76]$<br>$d = -2.62 [-3.74, -1.51]$<br>$d = -0.87 [-1.77, 0.03]$ |
| Avoidance acquisition | Freezing | Repeated measures ANOVA | $F(2, 30) = 1.60, p = 0.218$ | $\omega^2 = 0.04 [0.00, 1.00]$ |
| Avoidance acquisition | Suppression of lever pressing | Friedman test | $\chi^2(2) = 11.49, p = 0.003$ | $W = 0.52 [0.42, 1.00]$ |
| Avoidance acquisition | Suppression of lever pressing | Pairwise comparison with Wilcoxon rank sum test with Bonferroni correction | Day 1–2: $V = 23, p = 0.413$<br>Day 1–10: $V = 0, p < 0.001$<br>Day 2–10 : $V = 6, p = 0.032$ | $r = -0.3 [-0.75, 0.34]$<br>$r = -1 [-1, -1]$<br>$r = -0.78 [-0.94, -0.36]$ |
| Generalization | Avoidance | Mixed ANOVA | Stimuli: $F(2, 60) = 5.2, p = 0.008$<br>Day: $F(1, 60) = 0.29, p = 0.589$<br>S*D: $F(2, 60) < 0.01, p = 0.999$ | $\omega_p^2 = 0.11 [0.01, 1.00]$<br>$\omega_p^2 = -0.01 [0.00, 1.00]$<br>$\omega_p^2 = -0.03 [0.00, 1.00]$ |
| Generalization | Avoidance | Parametric pairwise contrasts with Tukey correction | CS–GS1: $t(60) = 0.29, p = 0.953$<br>CS–GS2: $t(60) = 2.93, p = 0.013$<br>GS1–GS2 : $t(60) = 2.63, p = 0.028$ | $d = 0.09 [-0.51, 0.9]$<br>$d = 0.88 [0.25, 1.51]$<br>$d = 0.79 [0.17, 1.41]$ |
| Generalization | Freezing | Non-parametric mixed ANOVA | Stimuli: $F(2, 50) = 13.02, p < 0.001$<br>Day: $F(1, 50) = 8.17, p = 0.006$<br>S*D: $F(2, 50) = 0.82, p = 0.444$ | $\omega_p^2 = 0.13 [0.02, 1.00]$<br>$\omega_p^2 = 0.04 [0.00, 1.00]$<br>$\omega_p^2 = -0.02 [0.00, 1.00]$ |
| Generalization | Freezing | Non-parametric pairwise contrasts with Tukey correction | CS–GS1: $t(50) = 1.02, p = 0.567$<br>CS–GS2: $t(50) = 4.84, p < 0.001$<br>GS1–GS2 : $t(50) = 3.82, p = 0.001$ | $d = 0.20 [-0.28, 0.68]$<br>$d = 0.95 [0.44, 1.46]$<br>$d = 0.75 [0.25, 1.25]$ |
| Generalization | Suppression of lever pressing | Mixed ANOVA | Stimuli: $F(2, 60) = 11.84, p < 0.001$<br>Day: $F(1, 60) = 4.2, p = 0.045$<br>S*D: $F(2, 60) = 0.05, p = 0.948$ | $\omega_p^2 = 0.25 [0.09, 1.00]$<br>$\omega_p^2 = 0.05 [0.00, 1.00]$<br>$\omega_p^2 = -0.03 [0.00, 1.00]$ |
| Generalization | Suppression of lever pressing | Parametric pairwise contrasts with Tukey correction | CS–GS1: $t(60) = 0.38, p = 0.92$<br>CS–GS2: $t(60) = 4.39, p < 0.001$ | $d = 0.12 [-0.49, 0.72]$<br>$d = 1.32 [0.67, 1.97]$ |
| | | | GS1–GS2 : $t(60) = 4.01$, **$p < 0.001$** | $d = 1.21$ [0.57, 1.85] |
| Extinction | Avoidance | Friedman test | $\chi^2(3) = 17.61$, **$p < 0.001$** | $W = 0.53$ [0.33, 1.00] |
| Extinction | Avoidance | Pairwise comparison with Wilcoxon rank sum test with Bonferroni correction | Day 1–2: $V = 49$, $p = 0.175$<br>Day 2–3: $V = 45$, $p = 0.306$<br>Day 3–4: $V = 51$, $p = 0.119$<br>Day 1–3: $V = 66$, **$p < 0.001$**<br>Day 1–4: $V = 65$, **$p = 0.002$**<br>Day 2–4: $V = 45$, **$p = 0.009$** | $r = 0.48$ [-0.14, 0.83]<br>$r = 0.36$ [-0.34, 0.81]<br>$r = 0.55$ [-0.09, 0.86]<br>$r = 1$ [1, 1]<br>$r = 0.97$ [0.89, 0.99]<br>$r = 1$ [1, 1] |
| Extinction | Freezing | Friedman test | $\chi^2(3) = 21.27$, **$p < 0.001$** | $W = 0.64$ [0.54, 1.00] |
| Extinction | Freezing | Pairwise comparison with Wilcoxon rank sum test with Bonferroni correction | Day 1–2: $V = 55$, $p = 0.053$<br>Day 2–3: $V = 43$, $p = 0.126$<br>Day 3–4: $V = 53$, $p = 0.083$<br>Day 1–3: $V = 61$, **$p = 0.01$**<br>Day 1–4: $V = 66$, **$p = 0.001$**<br>Day 2–4: $V = 66$, **$p = 0.004$** | $r = 0.67$ [0.14, 0.9]<br>$r = 0.56$ [-0.03, 0.86]<br>$r = 0.61$ [0.03, 0.88]<br>$r = 0.85$ [0.52, 0.96]<br>$r = 1$ [1, 1]<br>$r = 1$ [1, 1] |
| Extinction | Suppression of lever pressing | Repeated measures ANOVA | $F(3, 40) = 5.32$, **$p = 0.003$** | $\omega^2 = 0.23$ [0.03, 1.00] |
| Extinction | Suppression of lever pressing | Pairwise comparison with t-test with Tukey correction | Day 1–2: $t(40) = 1.24$, $p = 0.608$<br>Day 1–3: $t(40) = 2.61$, $p = 0.06$<br>Day 1–4: $t(40) = 3.75$, **$p = 0.003$**<br>Day 2–3: $t(40) = 1.37$, $p = 0.525$<br>Day 2–4: $t(40) = 2.52$, $p = 0.072$<br>Day 3–4: $t(40) = 1.15$, $p = 0.663$ | $d = 0.53$ [-0.34, 1.4]<br>$d = 1.11$ [0.21, 2.01]<br>$d = 1.6$ [0.67, 2.54]<br>$d = 0.58$ [-0.29, 1.46]<br>$d = 1.07$ [0.18, 1.97]<br>$d = 0.49$ [-0.38, 1.36] |

Bold values indicate statistical significance ( $p < 0.05$ ).

During the two-day generalization phase, female rats showed significant differences in avoidance behavior per stimulus (F(2, 60) = 5.2, p = 0.008, ω*_p_*^2^ = 0.11). While rats generalized their avoidance responses from the CS to GS1 (t(60) = 0.29, p = 0.953), they avoided more during CS than during GS2 presentations (t(60) = 2.93, p = 0.013, d = 0.88) and during GS1 than GS2 presentations (t(60) = 2.63, p = 0.028, d = 0.79). Freezing did significantly decrease from generalization day 1 to 2 (F(1, 50) = 8.17, p = 0.006, ω*_p_*^2^ = 0.04), but still differed between stimuli (F(2, 50) = 13.02, p < 0.001, ω*_p_*^2^ = 0.13). Again, rats froze more to the CS than during GS2 (t(50) = 4.84, p < 0.001, d = 0.95), and more during GS1 than GS2 (t(50) = 3.82, p = 0.001, d = 0.75), but showed similar levels of freezing to CS and GS1 (t(50) = 1.02, p = 0.567). The same pattern of results was found for suppression of lever pressing. Suppression decreased significantly from generalization day 1 to 2 (F(1, 60) = 4.2, p = 0.045, ω*_p_*^2^ = 0.05), and differed between stimuli (F(2, 60) = 11.84, p < 0.001, ω*_p_*^2^ = 0.25). Again, rats showed more suppression during the CS than during GS2 (t(60) = 4.39, p < 0.001, d = 1.32), and more suppression during GS1 than GS2 (t(60) = 4.01, p < 0.001, d = 1.21), but showed similar suppression of lever pressing during CS and GS1 (t(60) = 0.38, p = 0.92, d = 0.12). Essentially, we observed the same generalization gradient for the three defensive behaviors. Rats showed different responding for CS and GS2 presentations in avoidance, freezing and suppression of lever pressing and intermediate responding to GS1. Therefore, we obtained a successful generalization gradient in female rats.

Rats successfully extinguished avoidance (χ^2^ (3) = 17.61, p < 0.001, W = 0.53), freezing (χ^2^ (3) = 21.27, p < 0.001, W = 0.64) and suppression of lever pressing (F(3, 40) = 5.32, p = 0.003, ω2 = 0.23) over the course of the four extinction sessions. As in Experiment 1, avoidance and freezing were significantly decreased by the third session (avoidance: V = 66, p < 0.001, r = 1; freezing: V = 61, p = 0.01, r = 0.85). Suppression of lever pressing only reached statistically significant extinction by the fourth session (t(44) = 3.75, p = 0.003, d = 1.6).

For exploratory purposes, we compared the results in male rats (Experiment 1) to those obtained in female rats (Experiment 2). To summarize, when comparing both experiments, we found that male rats showed higher freezing during acquisition than female rats (F(1, 21) = 20.81, p < 0.001). Moreover, during generalization, interactions between sex and day were observed in avoidance (F(1, 105) = 6.83, p = 0.01, ω*_p_*^2^ = 0.01) and freezing (F(1, 105) = 32.02, p <0.001, ω*_p_*^2^ = 0.08). We observed that males showed a significant reduction of avoidance from generalization day 1 to 2 (V = 548, p < 0.001, r = 0.74), while this decrease in avoidance was not significant in females (V = 309, p = 0.405, r = 0.17). In contrast, both males (V = 666, p < 0.001, r = 0.54) and females (V = 433, p = 0.005, r = 0.54) showed a significant decrease in freezing over generalization tests. In suppression of lever pressing, we detected an interaction of sex and stimulus (F(2, 105) = 5.07, p = 0.008, ω _p_^2^ = 0.03).

While females suppressed lever pressing differently depending on the stimulus presented (Χ^2^(2) = 16.25, p<0.001, Ε^2^ = 0.25), males did not (Χ^2^(2) = 4.03, p = 0.133, Ε^2^ = 0.06). Detailed results can be found in Supplementary Table 3 and Supplementary Figure 5. To more systematically evaluate those apparent sex differences in the acquisition and generalization of platform-mediated avoidance, we conducted Experiment 3.

### Subtle sex differences in the modified version of the platform-mediated avoidance task

Here, we set out to replicate and confirm the sex differences observed during acquisition and generalization sessions in avoidance, freezing and suppression in the modified platform-mediated avoidance task that we observed across the previous two experiments, testing preregistered hypotheses derived from those preceding experiments (Figure 3), while also taking into account the females’ estrous stage.

**Figure 3.**
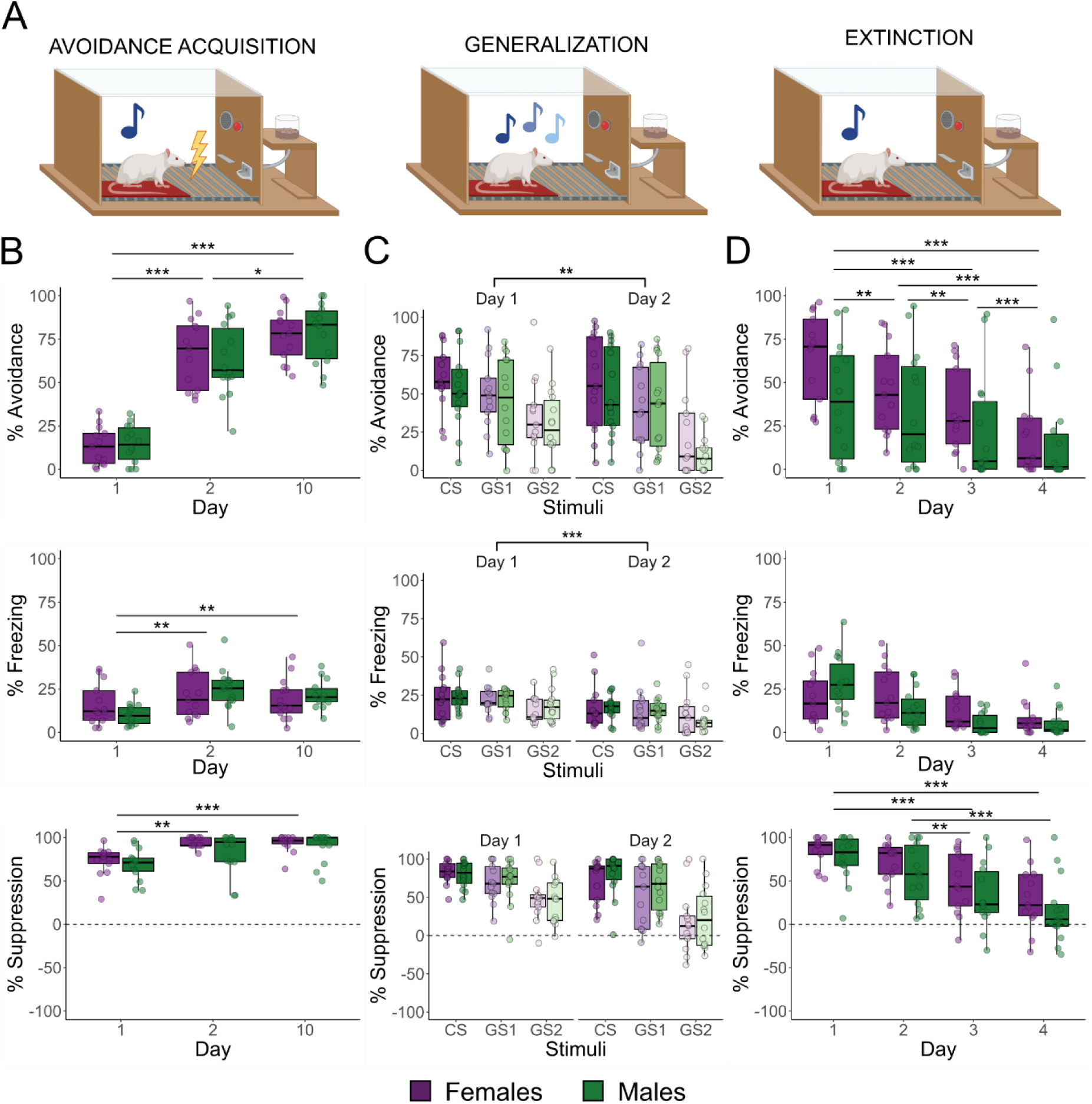
The boxplots represent the average of the first 3 CSs on a given day, except in the generalization test where they represent the average of each stimulus type on a given day. Results are expressed in % of tone duration. **A.** Graphic representation of the avoidance acquisition, generalization, and extinction phases. Different shades of blue denote different tone frequencies in the generalization test. **B.** Defensive behavior increased significantly over acquisition sessions in both sexes: avoidance (F(2, 75) = 114.21, p < 0.001), freezing (F(2, 50) = 8.63, p < 0.001) and suppression of lever pressing (F(2, 50) = 18.52, p <0.001). **C.** Both sexes showed a comparable decrease in avoidance behavior between generalization sessions 1 and 2 (F(1, 125) = 10.85, p = 0.001) and differences in avoidance between stimuli (F(2,125) = 51.18, p < 0.001). Rats avoided significantly more during the CS than during GS2 (V = 898, p = 0.004) and avoided more during GS1 than GS2 (V = 963, p < 0.001). We observed a decrease in freezing from generalization day 1 to 2 (F(1, 125) = 55.27, p < 0.001) and significant differences in freezing between stimuli (F(2,125) = 19.13, p < 0.001). For freezing, all pairwise comparisons between CS, GS1 and GS2 were significant (p < 0.001). For suppression of lever pressing during generalization, we found a significant interaction between day and stimulus (F(2, 125) = 4.34, p = 0.015), with specifically suppression of lever pressing during GS2 decreasing from session 1 to session 2 (χ^2^(1)= 7.61, p = 0.006). **D.** Extinction training significantly reduced avoidance (F(3,75) = 22.19, p < 0.001) and suppression of lever pressing (F(3,100) = 16.75, p < 0.001). We found an interaction between sex and day in freezing behavior (F(3, 75) = 4.41, p = 0.006). Female rats showed significantly more suppression than male rats (F(1, 100) = 4.44, p = 0.038).

Male and female rats successfully and similarly acquired avoidance behavior over the 10-day acquisition period (F(2, 75) = 114.21, p < 0.001, ω*_p_*^2^ = 0.74, see Table 3 for complete statistics). Avoidance behavior increased significantly from the first to the second session of avoidance acquisition (t(75) = –11.35, p < 0.001, d = –3.09) and was maintained over the course of training (Day 1-10: t(75) = –14.31, p < 0.001, d = –3.9; Day 2-10: t(75) = –2.96, p = 0.011, d = –0.81). Freezing levels (F(2, 50) = 8.63, p < 0.001, ω*_p_*^2^ = 0.12) and suppression of lever pressing (F(2, 50) = 18.52, p <0.001, ω*_p_*^2^ = 0.22) increased across training sessions and did not differ between sexes either.

**Table 3.** Statistical test results for Experiment 3.

| Test phase | Measured outcome | Statistical Test | Result | Effect size |
| --- | --- | --- | --- | --- |
| Avoidance acquisition | Avoidance | Mixed ANOVA | Sex: $F(1, 75) = 0.02, p = 0.878$<br>Day: $F(2, 75) = 114.21, p < 0.001$<br>S*D: $F(2, 75) = 0.24, p = 0.786$ | $\omega_p^2 = -0.01 [0, 1]$<br>$\omega_p^2 = 0.74 [0.65, 1]$<br>$\omega_p^2 = -0.02 [0, 1]$ |
| Avoidance acquisition | Avoidance | Pairwise comparison with t-test with Tukey correction | Day 1-2: $t(75) = -11.35, p < 0.001$<br>Day 1-10: $t(75) = -14.31, p < 0.001$<br>Day 2-10 : $t(75) = -2.96, p = 0.011$ | $d = -3.09 [-3.83, -2.35]$<br>$d = -3.9 [-4.73, -3.06]$<br>$d = -0.81 [-1.37, -0.25]$ |
| Avoidance acquisition | Freezing | Non-parametric mixed ANOVA | Sex: $F(1, 25) = 0.21, p = 0.649$<br>Day: $F(2, 50) = 8.63, p < 0.001$<br>S*D: $F(2, 50) = 2.04, p = 0.14$ | $\omega_p^2 = -0.01 [0, 1]$<br>$\omega_p^2 = 0.12 [0.02, 1]$<br>$\omega_p^2 = 0.01 [0, 1]$ |
| Avoidance acquisition | Freezing | Pairwise comparison with Wilcoxon rank sum test with Bonferroni correction | Day 1-2: $V = 59, p = 0.001$<br>Day 1-10: $V = 75, p = 0.005$<br>Day 2-10: $V = 225, p = 0.4$ | $r = -0.69 [-0.86, -0.39]$<br>$r = -0.6 [-0.81, -0.26]$<br>$r = 0.19 [-0.23, 0.55]$ |
| Avoidance acquisition | Suppression | Non-parametric mixed ANOVA | Sex: $F(1, 25) = 1.46, p = 0.239$<br>Day: $F(2, 50) = 18.52, p < 0.001$<br>S*D: $F(2, 50) = 0.66, p = 0.52$ | $\omega_p^2 = 0.03 [0, 1]$<br>$\omega_p^2 = 0.22 [0.09, 1]$<br>$\omega_p^2 = -0.01 [0, 1]$ |
| Avoidance acquisition | Suppression | Pairwise comparison with Wilcoxon rank sum test with Bonferroni correction | Day 1-2: $V = 65, p = 0.002$<br>Day 1-10: $V = 29, p < 0.001$<br>Day 2-10: $V = 105, p = 0.323$ | $r = -0.66 [-0.84, -0.34]$<br>$r = -0.85 [-0.93, -0.67]$<br>$r = -0.24 [-0.59, 0.19]$ |
| Generalization | Avoidance | Non-parametric mixed ANOVA | Sex: $F(1, 25) = 0.42, p = 0.523$<br>Day: $F(1, 125) = 10.84, p = 0.001$<br>Stimuli: $F(2, 125) = 51.18, p < 0.001$<br>Se*D: $F(1, 125) < 0.01, p = 0.98$<br>Se*St: $F(2, 125) = 0.43, p = 0.648$<br>D*St: $F(2, 125) = 2.14, p = 0.122$<br>Se*D*St: $F(2, 125) = 0.38, p = 0.686$ | $\omega_p^2 = 0.01 [0, 1]$<br>$\omega_p^2 = 0.02 [0, 1]$<br>$\omega_p^2 = 0.19 [0.1, 1]$<br>$\omega_p^2 = -0.01 [0, 1]$<br>$\omega_p^2 = -0.01 [0, 1]$<br>$\omega_p^2 < -0.01 [0, 1]$<br>$\omega_p^2 = -0.01 [0, 1]$ |
| Generalization | Avoidance | Non-parametric pairwise contrasts with Tukey correction | CS-GS1: $V = 893, p = 0.196$<br>CS-GS2: $V = 898, p = 0.004$<br>GS1-GS2: $V = 963, p < 0.001$ | $r = 0.2 [-0.1, 0.47]$<br>$r = 0.47 [0.18, 0.68]$<br>$r = 0.57 [0.31, 0.75]$ |
| Generalization | Freezing | Non-parametric mixed ANOVA | Sex: $F(1, 25) = 0.24, p = 0.63$<br>Day: $F(1, 125) = 55.27, p < 0.001$<br>Stimuli: $F(2, 125) = 19.13, p < 0.001$<br>Se*D: $F(1, 125) = 0.31, p = 0.58$<br>Se*St: $F(2, 125) = 0.41, p = 0.662$<br>D*St: $F(2, 125) = 0.25, p = 0.777$<br>Se*D*St: $F(2, 125) = 1.43, p = 0.243$ | $\omega_p^2 = -0.01 [0, 1]$<br>$\omega_p^2 = 0.08 [0.02, 1]$<br>$\omega_p^2 = 0.05 [0.01, 1]$<br>$\omega_p^2 < -0.01 [0, 1]$<br>$\omega_p^2 = -0.01 [0, 1]$<br>$\omega_p^2 = -0.01 [0, 1]$<br>$\omega_p^2 < -0.01 [0, 1]$ |
| Generalization | Freezing | Non-parametric pairwise contrasts with Tukey correction | CS-GS1: $V = 1258, p < 0.001$<br>CS-GS2: $V = 1226, p < 0.001$<br>GS1 – GS2: $V = 1250, p < 0.001$ | $r = 0.69 [0.5, 0.82]$<br>$r = 0.65 [0.44, 0.79]$<br>$r = 0.68 [0.48, 0.82]$ |
| Generalization | Suppression of lever pressing | Non-parametric mixed ANOVA | Sex: $F(1, 25) = 0.4, p = 0.53$<br>Day: $F(1, 125) = 11.69, p < 0.001$<br>Stimuli: $F(2, 125) = 49.01, p < 0.001$<br>Se*D: $F(1, 125) = 1.72, p = 0.191$<br>Se*St: $F(2, 125) = 0.07, p = 0.934$<br>D*St: $F(2, 125) = 4.34, p = 0.015$<br>Se*D*St: $F(2, 125) = 0.03, p = 0.964$ | $\omega_p^2 < -0.01 [0, 1]$<br>$\omega_p^2 = 0.04 [0.01, 1]$<br>$\omega_p^2 = 0.27 [0.17, 1]$<br>$\omega_p^2 < 0.01 [0, 1]$<br>$\omega_p^2 = -0.01 [0, 1]$<br>$\omega_p^2 = 0.01 [0, 1]$<br>$\omega_p^2 = -0.01 [0, 1]$ |
| Generalization | Suppression of lever pressing | Simple effects by stimuli: Krustal-Wallis test | CS - Day: $X^2(1) < 0.01, p = 0.965$<br>GS1 - Day: $X^2(1) = 0.68, p = 0.41$<br>GS2 - Day: $X^2(1) = 7.61, p = 0.006$ | $E^2_R < 0.01 [0, 1]$<br>$E^2_R = 0.01 [0, 1]$<br>$E^2_R = 0.14 [0.03, 1]$ |
| Extinction | Avoidance | Non-parametric mixed ANOVA | Sex: $F(1, 25) = 3.13, p = 0.089$<br>Day: $F(3, 75) = 22.19, p < 0.001$<br>S*D: $F(3, 75) = 1.60, p = 0.196$ | $\omega_p^2 = 0.04 [0, 1]$<br>$\omega_p^2 = 0.15 [0.04, 1]$<br>$\omega_p^2 = -0.01 [0, 1]$ |
| Extinction | Avoidance | Pairwise comparison with Wilcoxon rank sum test with Bonferroni correction | Day 1-2: $V = 277, p = 0.002$<br>Day 2-3: $V = 261, p = 0.008$<br>Day 3-4: $V = 247, p < 0.001$<br>Day 1-3: $V = 298, p < 0.001$<br>Day 1-4: $V = 307, p < 0.001$<br>Day 2-4: $V = 291, p < 0.001$ | $r = 0.7 [0.4, 0.87]$<br>$r = 0.61 [0.24, 0.82]$<br>$r = 0.79 [0.52, 0.92]$<br>$r = 0.83 [0.63, 0.93]$<br>$r = 0.89 [0.74, 0.95]$<br>$r = 0.79 [0.55, 0.91]$ |
| Extinction | Freezing | Non-parametric mixed ANOVA | Sex: $F(1, 25) = 0.07, p = 0.794$<br>Day: $F(3, 75) = 29.63, p < 0.001$<br>S*D: $F(3, 75) = 4.41, p = 0.006$ | $\omega_p^2 < 0.01 [0, 1]$<br>$\omega_p^2 = 0.23 [0.11, 1]$<br>$\omega_p^2 = 0.05 [0, 1]$ |
| Extinction | Freezing | Simple effects by sex: Krustal-Wallis test | Females: $X^2(3) = 9.94, p = 0.019$<br>Males: $X^2(3) = 24.17, p < 0.001$ | $E^2_R = 0.19 [0.08, 1]$<br>$E^2_R = 0.44 [0.29, 1]$ |
| Extinction | Suppression | Mixed ANOVA | Sex: $F(1, 100) = 4.44, p = 0.038$ | $\omega_p^2 = 0.03 [0, 1]$ |
| | | | Day: $F(3,100) = 16.75$, **$p < 0.001$**<br>S*D: $F(3, 100) = 0.06$, $p = 0.979$ | $\omega_p^2 = 0.3$ [0.17, 1]<br>$\omega_p^2 = -0.03$ [0, 1] |
| Extinction | Suppression | Pairwise comparison with t-test with Tukey correction | Day 1-2: $t(100) = 1.75$, $p = 0.31$<br>Day 1-3: $t(100) = 4.53$, **$p = 0.001$**<br>Day 1-4: $t(100) = 6.5$, **$p < 0.001$**<br>Day 2-3: $t(100) = 2.78$, **$p = 0.032$**<br>Day 2-4: $t(100) = 4.75$, **$p < 0.001$**<br>Day 3-4: $t(100) = 1.97$, $p = 0.205$ | $d = 0.48$ [-0.07, 1.02]<br>$d = 1.23$ [0.67, 1.8]<br>$d = 1.77$ [1.17, 2.37]<br>$d = 0.76$ [0.21, 1.31]<br>$d = 1.29$ [0.72, 1.87]<br>$d = 0.54$ [-0.01, 1.08] |

Bold values indicate statistical significance ( $p < 0.05$ ).

We observed no sex differences in avoidance, freezing, or suppression of lever pressing during the generalization sessions either. There was a reduction in avoidance between both sessions (F(1, 125) = 0.11, p = 0.001, ω*_p_*^2^ = 0.02) and a generalization gradient in the expected direction (F(2, 125) = 51.18, p < 0.001, ω*_p_*^2^ = 0.19). Specifically, no significant differences were present when comparing avoidance behavior during CS and GS1 (V = 893, p = 0.196), but avoidance differed between CS and GS2 (V = 898, p = 0.004, r = 0.47) and between GS1 and GS2 (V = 963, p < 0.001, r = 0.57). Similarly, we observed a reduction in freezing from generalization session 1 to 2 (F(1, 125) = 55.27, p < 0.001, ω*_p_*^2^ = 0.08) and pairwise differences in freezing between all stimuli (F(2, 125) = 19.13, p < 0.001, ω*_p_*^2^ = 0.05; CS-GS1: V = 1258, p < 0.001, r = 0.69; CS-GS2: V = 1226, p < 0.001, r = 0.65; GS1 – GS2: V = 1250, p < 0.001, r = 0.68), meaning that rats showed different freezing behavior for the different tones. For suppression of lever pressing, we found a significant interaction between day and stimulus type (F(2, 125) = 4.34, p = 0.015), with a reduction in suppression of lever pressing between both generalization sessions for GS2 only (χ^2^(1)= 7.61, p = 0.006, Ε^2^ = 0.14).

Four extinction sessions followed the generalization phase. Avoidance behavior extinguished significantly over the extinction sessions (F(3, 75) = 22.19, p < 0.001, ω*_p_*^2^ = 0.15), with a significant decrease already from the first to the second extinction session (V = 277, p = 0.002, r = 0.7), without any sex differences. We observed a significant interaction between sex and day for freezing during the extinction sessions (F(3, 75) = 4.41, p = 0.006, ω*_p_*^2^ = 0.05). Looking at simple effects of day per sex, we found that both females (Χ^2^(3)= 9.94, p = 0.019, Ε^2^ = 0.19) and males (Χ^2^(3)= 24.17, p < 0.001, Ε^2^ = 0.44) showed a reduction in freezing behavior over sessions. We also found a reduction in suppression of lever pressing over extinction sessions (F(3,100) = 16.75, p < 0.001, ω*_p_*^2^ = 0.3). Males showed significantly lower suppression of lever pressing than females (F(1, 100) = 4.44, p = 0.038). A significant decrease in suppression of lever pressing was apparent from the third session onwards (t(100) = 4.53, p < 0.001, d = 1.23).

Using DeepLabCut and SimBA, we processed our videos to obtain speed traces of each rat (see Methods and OSF for further information). We hypothesized that 40% of female rats would be classified as darters according to previous research [58]. However, in our case, 100% (13/13) of female rats were classified as darters and 86% (12/14) of male rats were also classified as darters. Due to the small numbers of non-darters, it is not possible to make reliable comparisons between darters and non-darters. Using a Chi-square test, we investigated whether there was a higher likelihood that females were classified as darters compared to males. However, the results did not support this exploratory hypothesis (χ^2^(1) = 0.4636, p = 0.496).

We assessed the estrous stage of the female rats for 10 days after the last extinction session and inferred estrous stage during training retrospectively to prevent a negative impact of the estrous staging procedure on our behavioral results. We divided the four stages of the estrous cycle into high estrogen phase (i.e., proestrus and estrus) and low estrogen phase (i.e., metestrus and diestrus) [59]. This categorization was determined by the estrous stage observed on the last day of extinction training. We then tested whether there were differences between the estrous groups for the three behavioral outcomes (avoidance, freezing, and suppression of lever pressing) on the last extinction day. We did not find significant differences in avoidance (t(11) = 1.37, p = 0.198, d = 0.76), freezing (t(11) = 1.16, p = 0.271) or suppression of lever pressing (t(11) = 1.27, p = 0.23, d = 0.68).

### Persistence of avoidance

In prior research using the platform-mediated avoidance task, persistent avoidance was reported in approximately 25% of (male) rats despite extinction [16] or extinction with response prevention [24]. Here, we used a preregistered criterion to classify a rat as a persistent avoider if on the first block of the first extinction day (i.e., after two sessions of generalization) its time on the platform during the CS presentation was more than 50%.

According to this definition, in Experiment 1, we found that 5/12 rats (42%) qualified as persistent avoiders. We analyzed differences between persistent avoiders and non-persistent avoiders in freezing and suppression of lever pressing on the last day of avoidance acquisition, but we did not find any significant differences between groups (see Supplementary Table S5 for all statistics). We also assessed differences in freezing and suppression of lever pressing between groups during the first extinction session. We did not find significant differences in freezing behavior, but we did find that persistent avoiders showed higher suppression of lever pressing than non-persistent avoiders (t(10) = –2.3, p = 0.044, d = –1.46).

In Experiment 2, 5 rats (5/11, 45%) met our predefined criterion for persistent avoidance. We conducted the same analyses as described for Experiment 1 and found no differences between the subgroups on the last avoidance acquisition session, nor on the first extinction day (see Supplementary Table S5 for all statistics).

In Experiment 3, 15 rats (15/27, 56%) met our preregistered criterion for persistent avoidance. We found no differences between persistent and non-persistent avoiders in terms of freezing or suppression of lever pressing on the last day of avoidance acquisition (see Supplementary Table S5). Freezing in the subgroups did not differ on the first day of extinction either. However, we did find a significant difference between persistent and non-persistent avoiders in suppression of lever pressing (F(1, 23) = 16.31, p < 0.001, ω*_p_*^2^ = 0.29), which is not surprising given that rats that avoided a lot during this first extinction day were classified as persistent avoiders, and avoidance implies that the rat is not able to press the lever. If we were to consider the first block of the third extinction session instead, as we have done previously [21], we would identify 6 persistent avoiders (6/27, 22.22%) which is consistent with previous literature [16,24]. On extinction day 1, we then find significantly more freezing (F(1, 23) = 8.49, p = 0.008, ω*_p_*^2^ = 0.22) and suppression of lever pressing (F(1, 23) = 5.87, p = 0.024, ω*_p_*^2^ = 0.09) in persistent avoiders than in non-persistent avoiders.

## Discussion

Here, we showed how avoidance, generalization, and extinction can be investigated in male and female rats, using a new, integrated platform-mediated avoidance task. Moreover, we developed scripts to facilitate the adoption of machine learning tools to analyze the animals’ behavior.

We first tested out the modified platform-mediated avoidance task in male rats (Experiment 1), and also validated it in female rats (Experiment 2). At the time of designing the experiments, only one published paper included male and female rats in a platform-mediated avoidance study in rats [33] and it did not include an extinction session, thus, it was important to test if our results would be comparable to Experiment 1. The combined findings from Experiment 1 and 2 indicated potential (if subtle) sex differences in avoidance, generalization, and extinction. To further explore these differences, we conducted Experiment 3, testing both sexes concurrently.

### Avoidance

Avoidance acquisition was successful in all three experiments, comparable to previous studies [14], particularly in the avoidance measurement and supported by freezing and suppression of lever pressing during the acquisition of avoidance. Females in Experiment 2 did not show an increase of freezing over acquisition sessions, unlike male rats in Experiment 1. This difference was not present in Experiment 3 where rats of both sexes showed similar freezing levels.

### Generalization

The generalization test phase inserted between avoidance acquisition and extinction yielded an orderly generalization gradient, where we observed markedly different responding between the CS and the more perceptually dissimilar GS2, with GS1 yielding intermediate responding. In Experiment 1, this generalization gradient was present for avoidance and suppression of lever pressing, while male rats showed similar levels of freezing across all stimuli. In contrast, in Experiment 2, females presented a generalization gradient across all measured behavioral outcomes. When comparing Experiment 1 and 2 we detected some sex differences. For instance, in males there was a reduction of avoidance behavior between generalization day 1 and 2 but this reduction in responding was not significant in females. In Experiment 3, where male and female rats were tested simultaneously, we did not find any sex differences in generalization.

### Extinction

After generalization testing, we performed extinction training. In Experiment 1 and 2 we observed a successful reduction of all fear-related behaviors, similar to other published studies [14]. Comparing both experiments, we did not observe any sex differences. Surprisingly, in Experiment 3 we did observe a significant interaction between sex and day for freezing over extinction. However, the effect was subtle, and both sexes showed a similar pattern of decreasing freezing during extinction. Additionally, females showed higher suppression of lever pressing than male rats.

### Sex differences

Across the presented studies, we observed minor sex differences only. The differences were more pronounced when comparing Experiments 1 and 2 than in Experiment 3, where subtle sex differences were observed in extinction only. The lack of expected sex differences in the third experiment, e.g., in generalization, may be explained by the increased experimental control afforded by testing both sexes concurrently. Additionally, Experiment 1 was performed by a male researcher and Experiment 2 by a female researcher, while in Experiment 3 both sexes were tested by one and the same female researcher (different from the researcher testing animals in Experiment 2). It has been reported previously in the literature that the sex of the experimenter can influence the measured outcomes of neuroscience experiments [60,61]. The absence of notable sex differences, even with the potential confounding factor of food motivation [62], is noteworthy. This suggests that the modified PMA protocol can be used to examine various manipulations and their impacts on avoidance, generalization, and extinction, with the assumption that any sex differences in a control condition will be subtle.

In adult rats, there have been no reported sex differences in acquisition of platform-mediated avoidance in Sprague-Dawley [33] or in Long-Evans [17]. However, the study in Long-Evans rats reported sex differences in extinction. They reported that male rats extinguished avoidance, but female rats did not exhibit a significant reduction in time spent on the platform over the extinction sessions. Additionally, they found that female rats showed higher freezing levels and less lever pressing during extinction. These results contradict our own. However, there are some differences to consider between both studies. First, we used a different rat strain. Second, Landin and Chandler [34] report a total number of lever pressing and lever pressing during tone presentation, whereas we report suppression of lever pressing. While these measures are related to each other, females could show less lever pressing but equal suppression of lever pressing to their male counterparts and vice versa. Lastly, their study design has some differences from ours. They had an avoidance test (but no generalization test) after acquisition, five sessions of reconditioning, and then five days of extinction training, which consisted of 15 CS presentations.

A recent study reported significant sex differences between male and female mice [22]. They showed that, after the 3^rd^ day of avoidance training, 48% of female mice were non-avoiders (receiving 3 or more shocks on the third training day) and that female avoiders and non-avoiders did not differ in their freezing during acquisition. After the third day, they performed an extinction training and a test session. They observed that females showed persistent avoidance after extinction, while males did not. These results are in contradiction to ours, but several important differences between both studies should be noted. Not only did they use mice, but their study left out the lever pressing component of the PMA task. While this may control for differences in food-seeking behavior between sexes [62], it removes the approach-avoidance conflict that makes the PMA procedure such a relevant translational tool to model clinical avoidance.

Sex differences have been reported in avoidance, generalization or extinction using other tasks than PMA. Studies using different active avoidance paradigms have reported that females acquire avoidance faster than males [38,63]. There have also been reports of sex differences in generalization testing. Females showed good discrimination after limited training, while males did not. However, after extended training, female rats began to generalize their fear responses due to impaired safety learning while male rats showed significant discrimination [64]. Another study found that females in low-estrogen phases of their estrous cycle showed discrimination, similar to males, while females in high-estrogen phases showed generalization between two different cues [65]. There is evidence suggesting that estrogen could be mediating effects of contextual fear generalization in a passive avoidance task [37]. Regarding extinction of avoidance responding, previous research reported that while both sexes showed similar extinction of avoidance, females showed stronger reductions in freezing than males [36]. There is more evidence of sex differences in fear extinction, in humans and in rodents, from hormonal effects [66,67] to neurobiological effects [68–70] (see review [71]). However, results in the literature are mixed and some authors did not observe sex differences in extinction in rats [72] or humans [73].

We investigated darting behavior in our third experiment, as previous research suggested that darting may be a conditioned behavior expressed largely by female rats [58,74], although further work challenged this finding in mice and rats [75,76]. Surprisingly, in our experiment we observed that all females were darters and 86% of male rats were considered darters as well. This was surprising because it was previously reported that the number of darters is around 40% in female rats and 10% in male rats [58]. The differences observed in this study compared to previous research may be due to previous research using Pavlovian fear conditioning only. In those experiments, darting is primarily an escape response. In contrast, in the present experiment, darting can serve either as a flight response towards a safe location (i.e., platform) or as an approach response to a reward (i.e., lever). Nevertheless, a previous experiment using PMA in mice found darters neither in males nor females [22].

### Persistent avoidance

Here, we used a preregistered criterion to classify a rat as a persistent avoider if on the first block of the first extinction day (i.e., after two sessions of generalization testing) its time on the platform during the CS presentation was more than 50%. While we were able to detect persistent avoiders (Experiment 1: 42%, Experiment 2: 45% and Experiment 3: 56%), we did not find consistent differences between persistent avoiders and non-persistent avoiders as reported in other studies [16,24]. This could have several reasons: (1) the introduction of a generalization phase in which the CS was partially reinforced, or (2) our use of a criterion to classify persistent avoiders that was not stringent enough to select true persistent avoider rats. To address the last possibility, we reanalyzed the data by applying criteria used in previous studies [16,24], but using those, we did not detect any persistent avoiders in Experiments 1 and 2. In Experiment 3, we found only 2 persistent avoiders (2/30: 7%), which is lower than the percentage found in those published studies and a value too small to permit robust analyses. If we considered the first block of the third extinction session instead in Experiment 3, as we have done previously [21], we find 6 persistent avoiders (6/27, 22.22%) which is consistent with previous literature [16,24]. We found that these persistent avoiders showed higher freezing and suppression of lever pressing than non-persistent avoiders on extinction day 1. However, these differences were not present at the end of acquisition as has been reported elsewhere [24].

### Open-source automated behavioral scoring in the PMA

We present a series of R scripts to facilitate the processing of outputs obtained from DeepLabCut and SimBA. The combined use of DeepLabCut and SimBA for the assessment of defensive behavior has already been validated by other authors [77]. These scripts are specifically adapted to analyze avoidance, freezing, and darting in the (modified) PMA task. Minimal coding knowledge is required, and we have provided explanations on OSF to help customize this solution to other interested users. These R scripts are readily adaptable to PMA tasks without generalization phase. The scripts are written to facilitate analysis of behaviors pre tone, during tone and post tone presentation. The use of supervised machine learning for the freezing analysis instead of purely pixel movement or motion indices [78], and our high ICC results for avoidance and freezing between a human scorer and the automated results, raise our confidence in this automated solution. Finally, sharing behavioral classifiers, analysis pipelines, and datasets obtained using such tools across laboratories could foster reproducibility in behavioral neuroscience [49].

#### Conclusions

In summary, this study presented a comprehensive behavioral procedure that can be used to study avoidance, generalization, and extinction in male and female rats. In addition, we provided a protocol including scripts to automatically analyze relevant behaviors in the platform-mediated avoidance setup using open-source tools.

## Declarations

### Ethics approval and consent to participate

All experiments were performed in accordance with Belgian and European laws (Belgian Royal Decree of 29/05/2013 and European Directive 2010/63/EU) and the ARRIVE 2.0 [40] guidelines and approved by the KU Leuven animal ethics committee (project license number: 011/2019).

### Consent for publication

Not applicable

### Availability of data and materials

The datasets generated and/or analyzed during the current study are available on the OSF repository: https://osf.io/ncb8j/.

### Author contributions

A.L.-M., L.L. and T.B. conceived and designed the study. A.L.-M. collected data and performed the data analysis. A.L.-M. wrote the first draft of the manuscript. L.L. and T.B. edited and critically reviewed the manuscript. All authors reviewed the results and approved the final version of the manuscript. L.L. and T.B. are co-last authors.

## Acknowledgements

We thank Nathan De Henau and Jialing Ding for assistance with data collection and behavioral scoring. We thank Celine Fatouh, Samantha Piers, Louca Bah D’Oliveira and Adrià Túnez Aquilué for assistance with behavioral scoring. We thank Mathijs Franssen for assistance with programming.

