## Supplemental figures and tables for "Generalization and extinction of platform-mediated avoidance in male and female rats"

**Supplementary Figure 1**

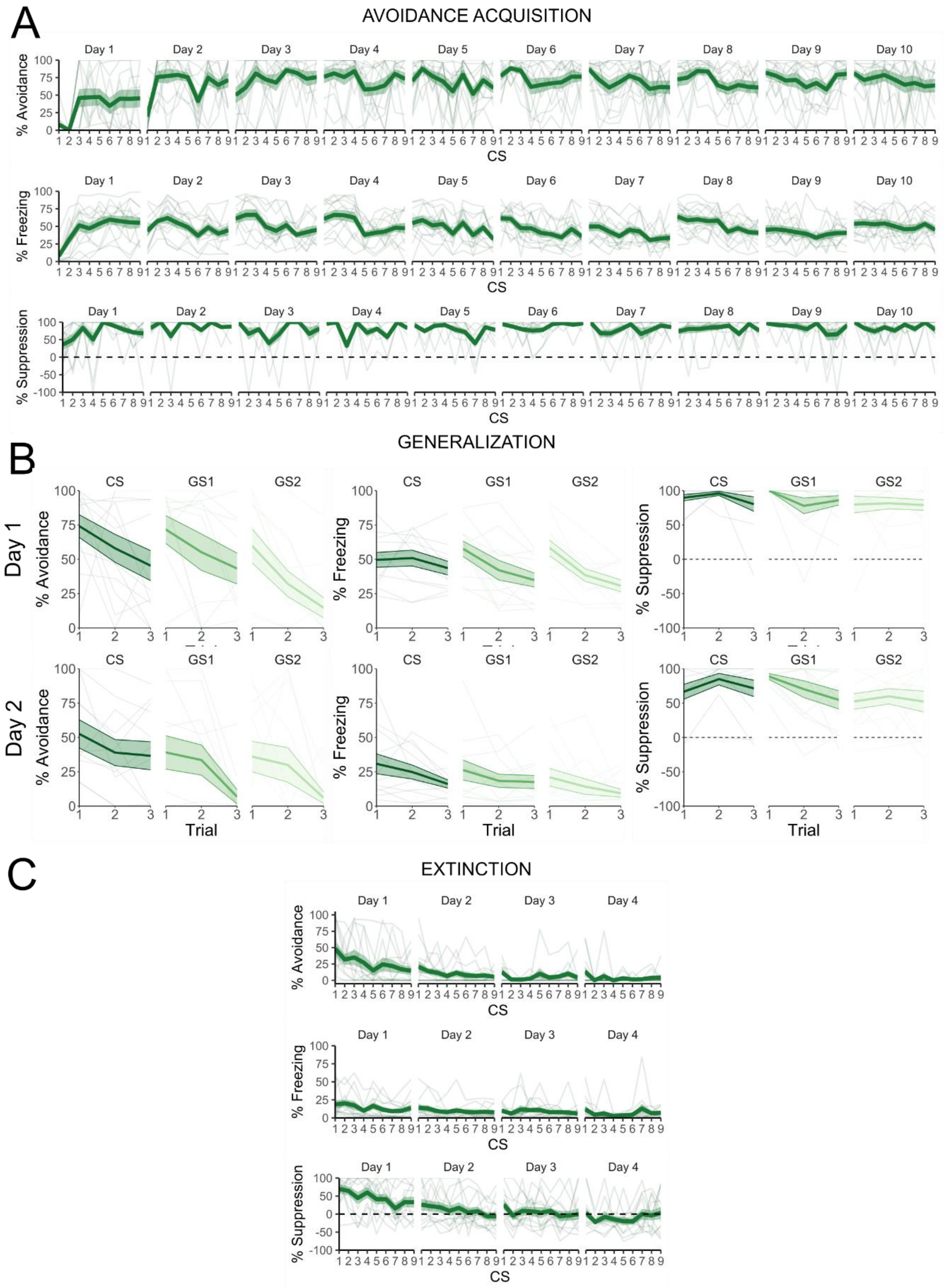


**Figure S1.** Trial-by-trial data of Experiment 1. The bold lines in the trial-by-trial plots represent the mean and the surrounding shaded area the standard error of the mean. Results are expressed in % of time of tone presentations. **A.** Avoidance, freezing, and suppression of lever pressing during acquisition training. **B.** Avoidance, freezing, and suppression of lever pressing during generalization testing. **C.** Avoidance, freezing, and suppression of lever pressing during extinction training.

**Supplementary Figure 2**

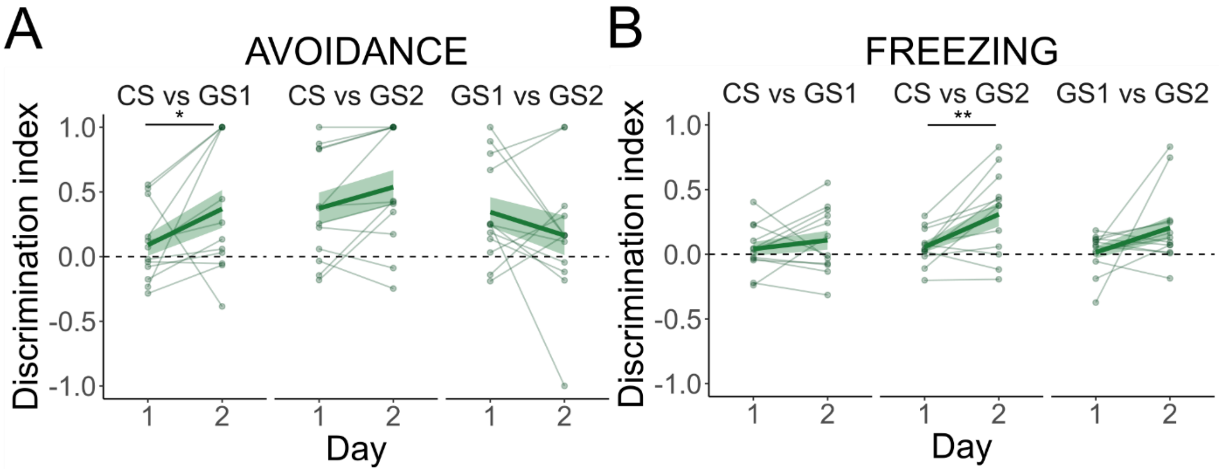


**Figure S2.** Discrimination indices during the generalization phase of Experiment 1. Discrimination indices were calculated by taking the difference between avoidance (or freezing) during the presentation of a given tone (e.g., CS) and another tone (e.g., GS1) and divided by the sum of avoidance or freezing to both tones. The bold lines represent the mean and the surrounding shaded area the standard error of the mean. Results are expressed as values ranging from -1 to 1. Values close to 1 or to -1 indicate high discrimination while values around 0 indicate low discrimination. **A.** Discrimination index of avoidance during generalization phase. There was a significant increase in the discrimination index between CS and GS1 from the first to the second session ($V = 12$, $p = 0.034$) **B.** Discrimination index of freezing during the generalization phase. There was a significant increase in the discrimination index between CS and GS2 from the first to the second generalization session ($t(11) = -3.17$, $p = 0.009$).

**Supplementary Figure 3**

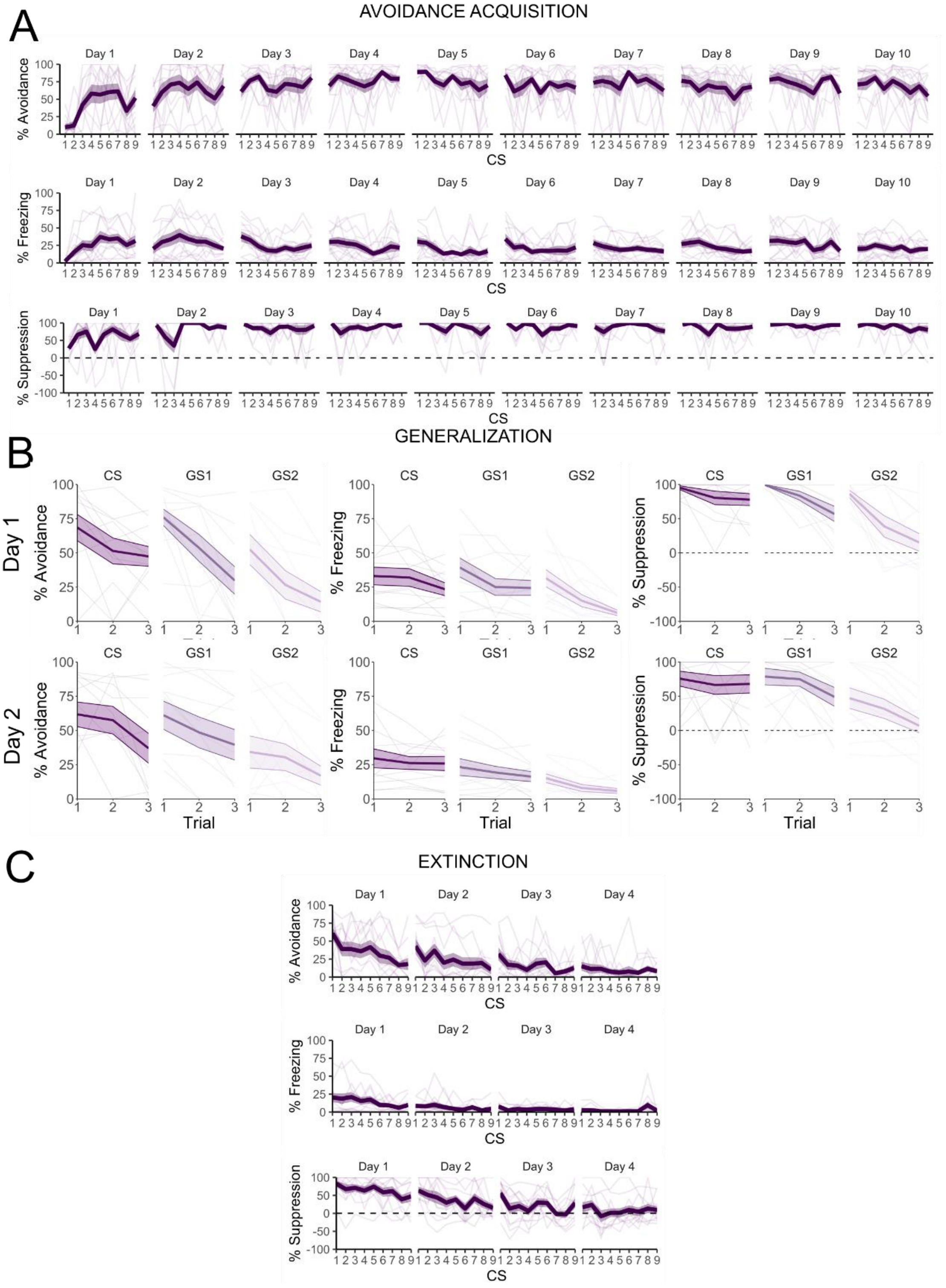


**Figure S3.** Trial-by-trial data of Experiment 2. The bold lines in the trial-by-trial plots represent the mean and the surrounding shaded area the standard error of the mean. Results are expressed in % of time of tone presentations. **A.** Avoidance, freezing, and suppression of lever pressing during the acquisition training phase. **B.** Avoidance, freezing, and suppression of lever pressing during the generalization testing. **C.** Avoidance, freezing, and suppression of lever pressing during the extinction training.

**Supplementary Figure 4**

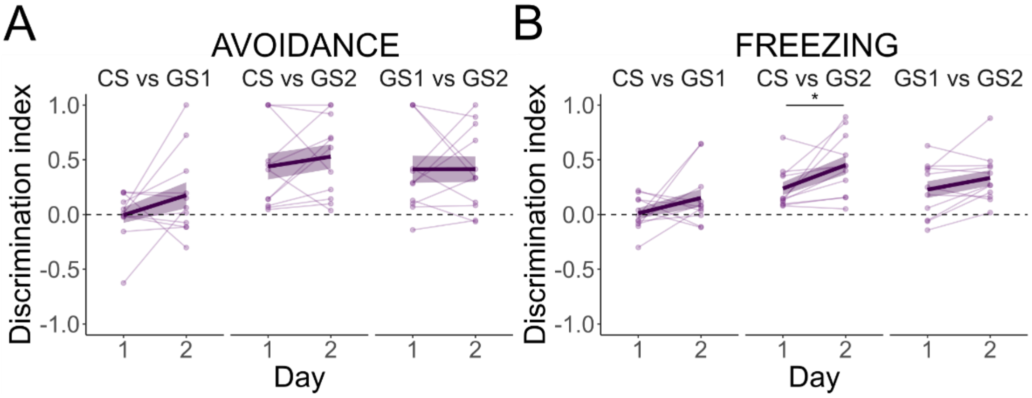


**Figure S4.** Discrimination indices during the generalization phase of Experiment 2. Discrimination indices were calculated by taking the difference between avoidance (or freezing) during the presentation of a given tone (e.g., CS) and another tone (e.g., GS1) and divided by the sum of avoidance or freezing to both tones. The bold lines represent the mean and the surrounding shaded area the standard error of the mean. Results are expressed as values ranging from -1 to 1. Values close to 1 or to -1 indicate high discrimination while values around 0 indicate low discrimination. **A.** Discrimination index of avoidance during the generalization phase. **B.** Discrimination index of freezing during the generalization phase. There was a significant increase in the discrimination index between CS and GS2 from the first to the second generalization session ($t(10) = -2.51$, $p = 0.031$).

**Supplementary Figure 5**

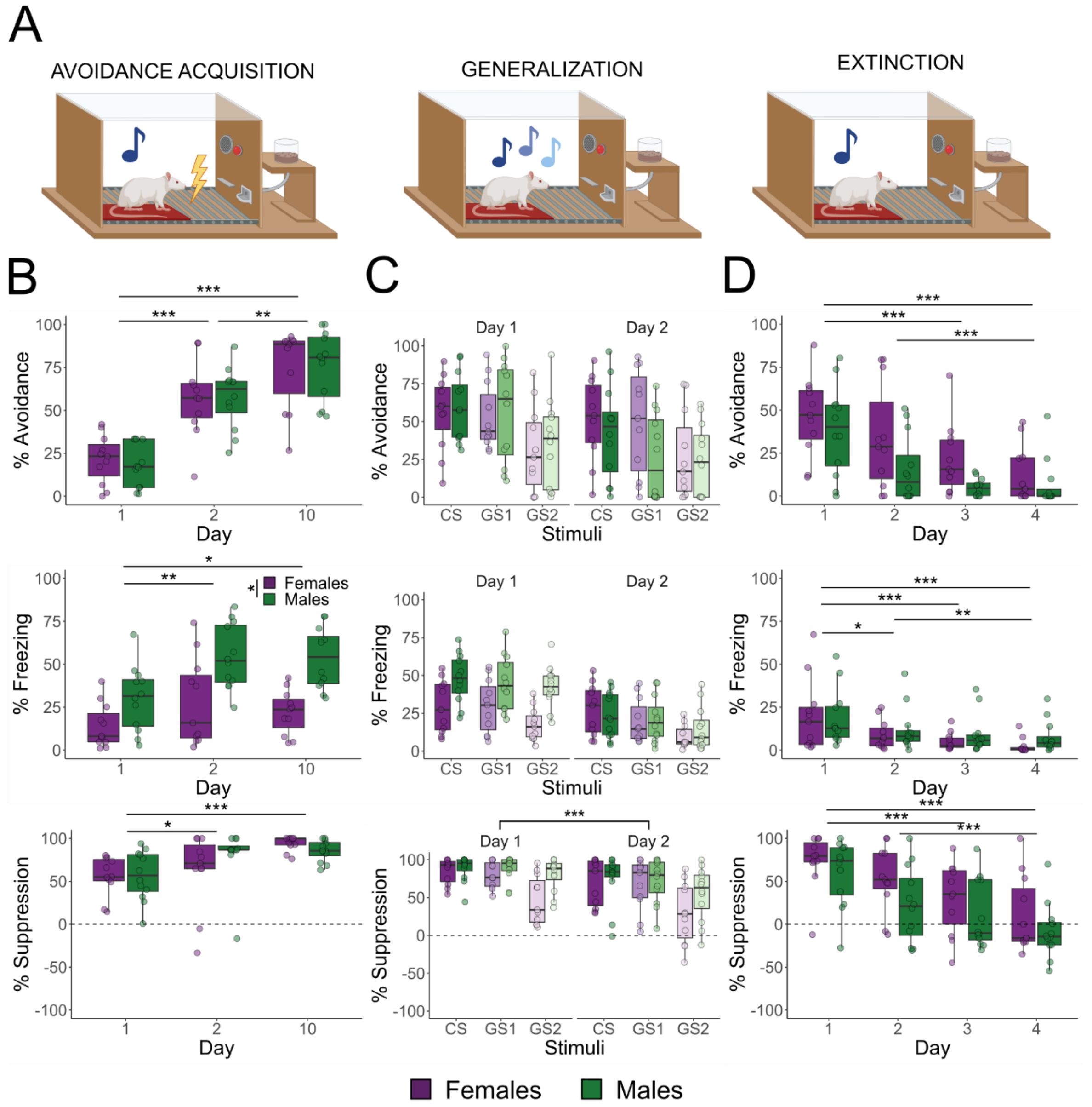


**Figure S5.** Results of Experiments 1 and 2 combined. The boxplots represent the average of the first 3 CSs on a given day, except in the generalization test where they represent the average of each stimulus type on a given day. Results are expressed in % of tone duration. **A.** Graphic representation of the avoidance acquisition, generalization and extinction phases. **B.** Rats of both sexes acquired avoidance similarly over acquisition sessions $F(2, 42) = 45.84$, $p < 0.001$). There was an overall increase in freezing ($F(2, 42) = 9.43$, $p < 0.001$), but this effect was driven by the males, and there were significant sex differences ($F(1, 21) = 20.81$, $p < 0.001$). Suppression of lever pressing increased significantly ($F(2, 42) = 14.94$, $p < 0.001$) over avoidance acquisition sessions for both sexes alike. **C.** There was a significant interaction between sex and generalization days for avoidance behavior ($F(1, 105) = 6.83$, $p = 0.01$). Investigating this interaction further, males showed a significant difference between days ($V = 548$, $p < 0.001$) but females did not. Regardless of sex, we found significant differences in avoidance to the different tones ($F(2, 105) = 21.63$, $p < 0.001$). Similarly, there was another interaction between sex and generalization day in freezing behavior ($F(1, 105) = 32.02$, $p < 0.001$). Investigating these effects by sex, we found a significant decrease in freezing from generalization day 1 to 2 in females ($V = 433$, $p = 0.005$) and in males ($V =$

666, p < 0.001). Again, there were significant differences between stimuli (F(2, 105) = 12.85, p <0.001). Regarding suppression of lever pressing, there was an interaction of sex by stimulus type. When assessing differences by sex, we observe that females, but not males, suppressed lever pressing differently depending on stimulus (F(2, 105) = 5.07, p = 0.008). In both sexes, there was a reduction in suppression of lever pressing from generalization day 1 to 2 (F(1, 105) = 14.06, p <0.001). **D.** Avoidance behavior decreased in both sexes across extinction sessions (F(3, 63) = 19.83, p <0.001). Likewise for freezing (F(3, 63) = 13.52, p <0.001) and suppression of lever pressing (F(3, 63) = 24.07, p <0.001)

**Supplementary Figure 6**

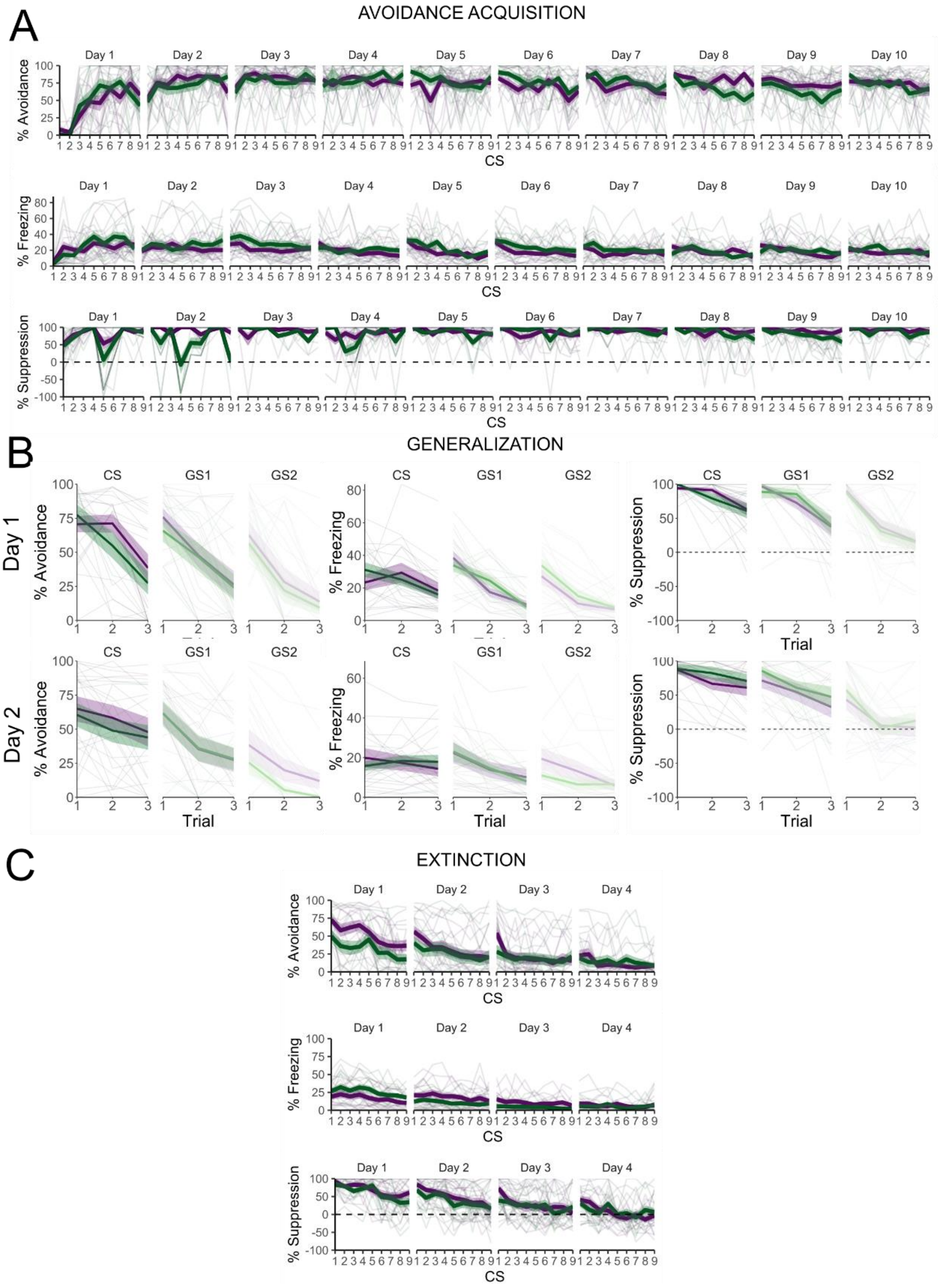


**Figure S6.** Trial-by-trial data of Experiment 3. The bold lines in the trial-by-trial plots represent the mean and the surrounding shaded area the standard error of the mean. Results are expressed in % of time during tone presentations. **A.** Avoidance, freezing, and suppression of lever pressing during the avoidance acquisition phase. **B.** Avoidance, freezing and suppression of lever pressing during the generalization phase. **C.** Avoidance, freezing and suppression of lever pressing during the extinction phase.

**Supplementary Figure 7**

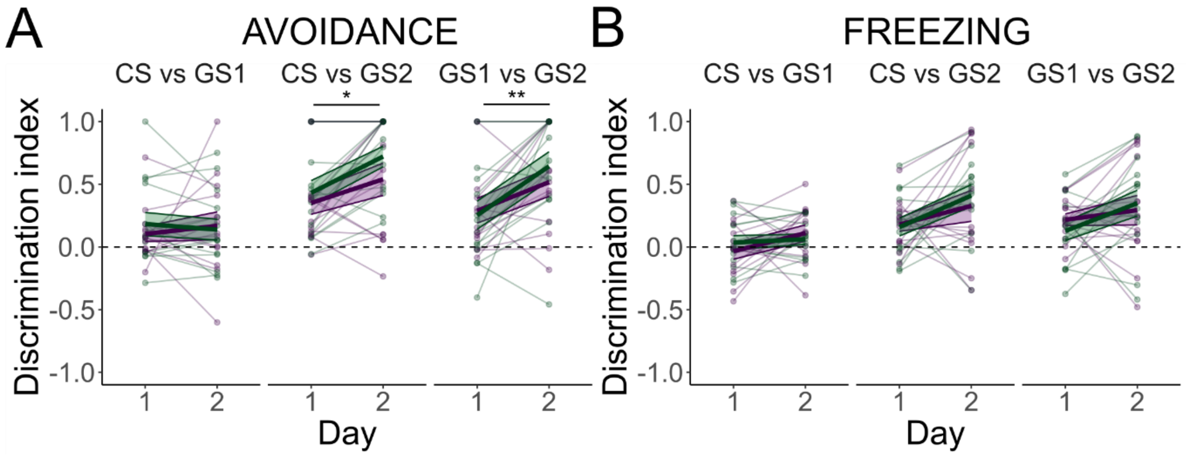


**Figure S7.** Discrimination indices during the generalization phase of Experiment 3. Discrimination indices were calculated by taking the difference between avoidance (or freezing) during the presentation of a given tone (e.g., CS) and another tone (e.g., GS1) and divided by the sum of avoidance (or freezing) to both tones. The bold lines represent the mean and the surrounding shaded area the standard error of the mean. Results are expressed as values ranging from -1 to 1. Values close to 1 or to -1 indicate high discrimination while values around 0 indicate low discrimination. **A.** Discrimination index of avoidance during generalization phase. There was a significant increase in discrimination index from day 1 to 2 in the CS and GS2 comparison ($F(1, 50) = 5.77$, $p = 0.02$) and in the GS1 and GS2 comparison ($F(1, 50) = 8.1$, $p = 0.006$). **B.** Discrimination index of freezing during the generalization phase.

Table S1. Discrimination indices in Experiment 1

| **Phase** | **Measured outcome** | **Statistical Test** | **Result** | **Effect size** |
|---|---|---|---|---|
| Generalization | Discrimination index avoidance CS – GS1 | Wilcox test | V = 12, **p = 0.034** | r = -0.69 [-0.90, -0.21] |
| Generalization | Discrimination index avoidance CS – GS2 | Wilcox test | V = 14, p = 0.1 | r = -0.58 [-0.86, -0.02] |
| Generalization | Discrimination index avoidance GS1 – GS2 | Wilcox test | V = 49, p = 0.4697 | r = 0.26 [-0.36, 0.72] |
| Generalization | Discrimination index freezing CS – GS1 | Paired t-test | t(11) = -0.89, p = 0.39 | d = -0.26, [-1.01, 0.31] |
| Generalization | Discrimination index freezing CS – GS2 | Paired t-test | t(11) = -3.17, **p = 0.009** | d = -0.92 [-1.69, -0.44] |
| Generalization | Discrimination index freezing GS1 – GS2 | Paired t-test | t(11) = -1.73, p = 0.11 | d = -0.5 [-0.97, -0.1] |

Table S2. Discrimination indices in Experiment 2

| **Phase** | **Measured outcome** | **Statistical Test** | **Result** | **Effect size** |
|---|---|---|---|---|
| Generalization | Discrimination index avoidance CS – GS1 | Paired t-test | t(10) = -1.39, p = 0.195 | d = -0.42 [-1.07, 0.17] |
| Generalization | Discrimination index avoidance CS – GS2 | Wilcox test | V = 19, p = 0.415 | r = -0.31 [-0.76, 0.34] |
| Generalization | Discrimination index avoidance GS1 – GS2 | Wilcox test | V = 33, p = 1 | r = 0 [-0.58, 0.58] |

| | | | | |
|---|---|---|---|---|
| Generalization | Discrimination index freezing CS – GS1 | Paired t-test | t(10) = -1.37, p = 0.201 | d = -0.41 [-1.02, 0.18] |
| Generalization | Discrimination index freezing CS – GS2 | Paired t-test | **t(10) = -2.51, p = 0.031** | d = -0.76 [-1.37, -0.36] |
| Generalization | Discrimination index freezing GS1 – GS2 | Paired t-test | t(10) = -1.83, p = 0.097 | d = -0.55 [-1.26, -0.06] |

Table S3. Comparison males in Experiment 1 and females in Experiment 2

| **Phase** | **Measured outcome** | **Statistical Test** | **Result** | **Effect size** |
|---|---|---|---|---|
| Avoidance acquisition | Avoidance | Mixed ANOVA | Sex: F(1, 21) = 0.02, p = 0.878<br>Day: F(2, 42) = 45.84, **p < 0.001**<br>S*D: F(2, 42) = 0.09, p = 0.915 | Sex: $\omega_p^2$ = -0.01 [0, 1]<br>Day: $\omega_p^2$ = 0.6 [0.47, 1]<br>S*D: $\omega_p^2$ = -0.03 [0, 1] |
| Avoidance acquisition | Avoidance | Pairwise comparisons using t-test with Tukey correction | Day 1-2: t(63) = -6.76, **p < 0.001**<br>Day 1-10: t(63) = -10.04, **p < 0.001**<br>Day 2-10: t(63) = -3.28, **p = 0.004** | Day 1-2: d = -2 [-2.68, -1.31]<br>Day 1-10: d = -2.96 [-3.76, -2.17]<br>Day 2-10: d = 0.97 [-1.58, -0.35 |
| Avoidance acquisition | Freezing | Mixed ANOVA | Sex: F(1, 21) = 20.81, **p < 0.001**<br>Day: F(2, 42) = 9.43, **p < 0.001**<br>S*D: F(2, 42) = 1.48, p = 0.239 | Sex: $\omega_p^2$ = 0.27 [0.13, 1]<br>Day: $\omega_p^2$ = 0.11 [0.01, 1]<br>S*D: $\omega_p^2$ = 0 [0, 1] |
| Avoidance acquisition | Freezing | Pairwise comparisons using t-test with Tukey correction | Day 1-2: t(63) = -3.51, **p = 0.002**<br>Day 1-10: t(63) = -2.9, **p = 0.014**<br>Day 2-10: t(63) = 0.6, p = 0.818 | Day 1-2: d = -1.04 [-1.65, -0.42]<br>Day 1-10: d = -0.86 [-1.47, -0.25]<br>Day 2-10: d = 0.18 [-0.41, 0.77] |
| Avoidance acquisition | Suppression of lever pressing | Non-parametric mixed ANOVA | Sex: F(1, 21) = 0.76, p =0.392<br>Day: F(2, 42) = 14.94, **p <0.001**<br>S*D: F(2, 42) = 2.85, p = 0.069 | Sex: $\omega_p^2$ = -0.01 [0, 1]<br>Day: $\omega_p^2$ = 0.18 [0.05, 1]<br>S*D: $\omega_p^2$ = 0.01 [0, 1] |
| Avoidance acquisition | Suppression of lever pressing | Wilcoxon signed ranks test | Day 1-2: V = 67, **p = 0.03**<br>Day 1-10: V = 13, **p < 0.001**<br>Day 2-10: V = 74, p = 0.154 | Day 1-2: r = -0.51 [-0.78, -0.1]<br>Day 1-10: r = -0.91 [-0.96, -0.78]<br>Day 2-10: r = -0.36 [-0.69, 0.09] |
| Generalization | Avoidance | Non-parametric mixed ANOVA | Sex: F(1, 21) = 0.18, p = 0.677<br>Day: F(1, 105) = 16.04, **p < 0.001**<br>Stimuli: F(2, 105) = 21.63, **p < 0.001** | Sex: $\omega_p^2$ = -02 [0, 1]<br>Day: $\omega_p^2$ = 0.04 [0, 1]<br>Stimuli: $\omega_p^2$ = 0.09 [0.02, 1] |

| | | | | |
|---|---|---|---|---|
| | | | Se*D: F(1, 105) = 6.83, **p = 0.01**<br>Se*St: F(2, 105) = 1.11, p = 0.335<br>D*St: F(2, 105) = 0.96, p = 0.386<br>Se*D*St: F(2, 105) = 0.91, p = 0.404 | Se*D: $\omega_p^2$ = 0.01 [0, 1]<br>Se*St: $\omega_p^2$ = 0 [0, 1]<br>D*St: $\omega_p^2$ = 0 [0, 1]<br>Se*D*St: $\omega_p^2$ = 0 [0, 1] |
| Generalization | Avoidance | Simple effects by sex; Wilcoxon signed ranks test | Females – Day: V = 309, p = 0.405<br>Males – Day: V = 548, **p < 0.001** | r = 0.17 [-0.23, 0.52]<br>r = 0.74 [0.52, 0.87] |
| Generalization | Avoidance | Wilcoxon signed ranks test with Bonferroni correction | CS-GS1: V = 812, **p = 0.002**<br>CS-GS2: V = 710, **p = 0.012**<br>GS1-GS2: V = 757, **p = 0.002** | CS-GS1: r = 0.51 [0.22, 0.71]<br>CS-GS2: r = 0.43 [0.12, 0.67]<br>GS1-GS2: r = 0.53 [0.24, 0.73] |
| Generalization | Freezing | Non-parametric mixed ANOVA | Sex: F(1, 21) = 3.64, p = 0.07<br>Day: F(1, 105) = 107.87, **p < 0.001**<br>Stimuli: F(2, 105) = 12.85, **p < 0.001**<br>Se*D: F(1, 105) = 32.02, **p < 0.001**<br>Se*St: F(2, 105) = 1.73, p = 0.18<br>D*St: F(2, 105) = 0.64, p = 0.53<br>Se*D*St: F(2, 105) = 0.75, p = 0.473 | Sex: $\omega_p^2$ = 0.1 [0.03, 1]<br>Day: $\omega_p^2$ = 0.23 [0.13, 1]<br>Stimuli: $\omega_p^2$ = 0.07 [0.01, 1]<br>Se*D: $\omega_p^2$ = 0.08 [0.02, 1]<br>Se*St: $\omega_p^2$ = 0 [0, 1]<br>D*St: $\omega_p^2$ = -0.01 [0, 1]<br>Se*D*St: $\omega_p^2$ = -0.01 [0, 1] |
| Generalization | Freezing | Simple effects by sex; Wilcoxon signed ranks test | Females – Day: V = 433, **p = 0.005**<br>Males – Day: V = 666, **p < 0.001** | r = 0.54 [0.21, 0.76]<br>r = 1 [1, 1] |
| Generalization | Freezing | Wilcoxon signed ranks test with Bonferroni correction | CS-GS1: V = 999, **p < 0.001**<br>CS-GS2: V = 1019, **p < 0.001**<br>GS1-GS2: V = 1038, **p < 0.001** | CS-GS1: r = 0.85 [0.73, 0.92]<br>CS-GS2: r = 0.89 [0.79, 0.94]<br>GS1-GS2: r = 0.92 [0.85, 0.96] |
| Generalization | Suppression of lever pressing | Non-parametric mixed ANOVA | Sex: F(1, 21) = 3.32, p = 0.082<br>Day: F(1, 105) = 14.06, **p < 0.001**<br>Stimuli: F(2, 105) = 17.07, **p < 0.001**<br>Se*D: F(1, 105) = 0.04, p = 0.844<br>Se*St: F(2, 105) = 5.07, **p = 0.008**<br>D*St: F(2, 105) = 1.23, p = 0.297<br>Se*D*St: F(2, 105) = 0.10, p = 0.9 | Sex: $\omega_p^2$ = 0.05 [0, 1]<br>Day: $\omega_p^2$ = 0.08 [0.02, 1]<br>Stimuli: $\omega_p^2$ = 0.14 [0.05, 1]<br>Se*D: $\omega_p^2$ = 0 [0, 1]<br>Se*St: $\omega_p^2$ = 0.03 [0, 1]<br>D*St: $\omega_p^2$ = -0.01 [0, 1]<br>Se*D*St: $\omega_p^2$ = -0.01 [0, 1] |
| Generalization | Suppression of lever pressing | Simple effects by sex; Kruskal-Wallis test | Females – Stimuli: $X^2(2)$ = 16.25, **p < 0.001**<br>Males – Stimuli: $X^2(2)$ = 4.03, p = 0.133 | Females – Stimuli: $E^2_R$= 0.25 [0.13, 1]<br>Males – Stimuli: $E^2_R$= 0.06 [0.01, 1] |

| | | | | |
|---|---|---|---|---|
| Extinction | Avoidance acquisition | Non-parametric mixed ANOVA | Sex: F(1, 21) = 3.36, p = 0.081<br>Day: F(3, 63) = 19.83, **p < 0.001**<br>S*D: F(3, 63) = 0.99, p = 0.402 | Sex: $\omega_p^2$ = 0.07 [0.01, 1]<br>Day: $\omega_p^2$ = 0.26 [0.12, 1]<br>S*D: $\omega_p^2$ = -0.02 [0, 1] |
| Extinction | Avoidance acquisition | Wilcoxon signed rank test with Bonferroni correction | Day 1-2: V = 214.5, **p = 0.004**<br>Day 2-3: V = 168, p = 0.071<br>Day 3-4: V = 146, p = 0.13<br>Day 1-3: V = 253, **p < 0.001**<br>Day 1-4: V = 267, **p < 0.001**<br>Day 2-4: V = 174, **p = 0.001** | Day 1-2: r = 0.7 [0.36, 0.87]<br>Day 2-3: r = 0.45 [-0.07, 0.78]<br>Day 3-4: r = 0.39 [-0.1, 0.73]<br>Day 1-3: r = 1 [1, 1]<br>Day 1-4: r = 0.93 [0.84, 0.97]<br>Day 2-4: r = 0.83 [0.56, 0.94] |
| Extinction | Freezing | Non-parametric mixed ANOVA | Sex: F(1, 21) = 0.1, p = 0.76<br>Day: F(3, 63) = 13.52, **p < 0.001**<br>S*D: F(3, 63) = 0.39, p = 0.758 | Sex: $\omega_p^2$ = 0 [0, 1]<br>Day: $\omega_p^2$ = 0.14 [0.03, 1]<br>S*D: $\omega_p^2$ = -0.02 [0, 1] |
| Extinction | Freezing | Wilcoxon signed rank test with Bonferroni correction | Day 1-2: V = 222, **p = 0.011**<br>Day 2-3: V = 178, p = 0.098<br>Day 3-4: V = 187, p = 0.14<br>Day 1-3: V = 251, **p < 0.001**<br>Day 1-4: V = 269, **p < 0.001**<br>Day 2-4: V = 239, **p = 0.002** | Day 1-2: r = 0.61 [0.24, 0.83]<br>Day 2-3: r = 0.41 [-0.03, 0.72]<br>Day 3-4: r = 0.36 [-0.1, 0.68]<br>Day 1-3: r = 0.82 [0.6, 0.92]<br>Day 1-4: r = 0.95 [0.88, 0.98]<br>Day 2-4: r = 0.73 [0.43, 0.89] |
| Extinction | Suppression of lever pressing | Mixed ANOVA | Sex: F(1, 21) = 2.6, p = 0.122<br>Day: F(3, 63) = 24.07, **p < 0.001**<br>S*D: F(3, 63) = 0.38, p = 0.77 | Sex: $\omega_p^2$ = 0.06 [0, 1]<br>Day: $\omega_p^2$ = 0.26 [0.12, 1]<br>S*D: $\omega_p^2$ = -0.03 [0, 1] |
| Extinction | Suppression of lever pressing | Pairwise comparisons using t-test with Tukey correction | Day 1-2: t(84) = 2.51, p = 0.66<br>Day 1-3: t(84) = 4.01, **p < 0.001**<br>Day 1-4: t(84) = 5.672, **p < 0.001**<br>Day 2-3: t(84) = 1.506, p = 0.439<br>Day 2-4: t(84) = 3.163, **p = 0.011**<br>Day 3-4: t(84) = 1.658, p = 0.352 | Day 1-2: d = 0.74 [0.14, 1.34]<br>Day 1-3: d = 1.18 [0.57, 1.8]<br>Day 1-4: d = 1.67 [1.03, 2.32]<br>Day 2-3: d = 0.44 [-0.15, 1.04]<br>Day 2-4: d = 0.93 [0.33, 1.54]<br>Day 3-4: d = 0.49 [-0.1, 1.08] |

Table S4. Discrimination indices in Experiment 3

| **Phase** | **Measured outcome** | **Statistical Test** | **Result** | **Effect size** |
|---|---|---|---|---|
| Generalization | Discrimination index avoidance CS – GS1 | Non-parametric mixed ANOVA | Day: F (1, 25) = 0.02, p = 0.884<br>Sex: F(1, 25) = 0.03, p = 0.086<br>S*D: F(1, 25) = 0.26, p = 0.616 | $\omega_p^2$ = -0.02 [0, 1]<br>$\omega_p^2$ = -0.02 [0, 1]<br>$\omega_p^2$ = -0.01 [0, 1] |

| | | | | |
|---|---|---|---|---|
| Generalization | Discrimination index avoidance CS – GS2 | Mixed ANOVA | Day: F (1, 50) = 5.77, **p = 0.02**<br>Sex: F(1, 50) = 1.7, p = 0.198<br>S*D: F(1, 50) = 0.25, p = 0.619 | $\omega_p^2$ = 0.08 [0, 1]<br>$\omega_p^2$ = 0.01 [0, 1]<br>$\omega_p^2$ = -0.01 [0, 1] |
| Generalization | Discrimination index avoidance GS1 – GS2 | Mixed ANOVA | Day: F (1, 50) = 8.1, **p = 0.006**<br>Sex: F(1, 50) = 0.16, p = 0.694<br>S*D: F(1, 50) = 0.5, p = 0.484 | $\omega_p^2$ = 0.12 [0.01, 1]<br>$\omega_p^2$ = -0.02 [0, 1]<br>$\omega_p^2$ = -0.01 [0, 1] |
| Generalization | Discrimination index freezing CS – GS1 | Mixed ANOVA | Day: F (1, 50) = 2.21, p = 0.143<br>Sex: F(1, 50) = 0.03, p = 0.86<br>S*D: F(1, 50) = 1.08, p = 0.304 | $\omega_p^2$ = 0.02 [0.01, 1]<br>$\omega_p^2$ = -0.02 [0, 1]<br>$\omega_p^2$ < 0.01 [0, 1] |
| Generalization | Discrimination index freezing CS – GS2 | Mixed ANOVA | Day: F (1, 50) = 5.35, **p = 0.025**<br>Sex: F(1, 50) = 0.12, p = 0.728<br>S*D: F(1, 50) = 0.34, p = 0.56 | $\omega_p^2$ = 0.07 [0, 1]<br>$\omega_p^2$ = -0.02 [0, 1]<br>$\omega_p^2$ = -0.01 [0, 1] |
| Generalization | Discrimination index freezing GS1 – GS2 | Repeated measures ANOVA | Day: F (1, 50) = 2.85 p = 0.097<br>Sex: F(1, 50) = 0.03, p = 0.851<br>S*D: F(1, 50) = 0.65, p = 0.423 | $\omega_p^2$ = 0.03 [0, 1]<br>$\omega_p^2$ = -0.02 [0, 1]<br>$\omega_p^2$ = -0.01 [0, 1] |

Table S5. Persistent avoiders compared to non-persistent avoiders.

| **Phase** | **Measured outcome** | **Statistical Test** | **Result** | **Effect size** |
|---|---|---|---|---|
| **Experiment 1** | | | | |
| Avoidance | Freezing | T-test | t(10) = -2.12, p = 0.06 | d = -1.25 [-4.49, 0.01] |
| Avoidance | Suppression of lever pressing | T-test | t(10) = -0.87, p = 0.4 | d = -0.54 [-2.34, 0.72] |
| Extinction | Freezing | Welch's t-test | t(4.8) = -1.78, p = 0.14 | d = -1.11 [-5.6, 0.01] |
| Extinction | Suppression of lever pressing | T-test | t(10) = -2.3, **p = 0.044** | d = -1.46 [-3.6, -0.65] |
| **Experiment 2** | | | | |
| Avoidance | Freezing | T-test | t(9) = 0.17, p = 0.87 | d = 0.1 [-1.48, 1.69] |
| Avoidance | Suppression of lever pressing | Welch's t-test | t(4.68) = 2.08, p = 0.096 | d = 1.31 [0.25, 4.53] |
| Extinction | Freezing | Welch's t-test | t(6.88) = 0.7, p = 0.5 | d = 0.41 [-1.62, 1.72] |
| Extinction | Suppression of lever pressing | Wilcoxon rank sum test | W = 6, p = 0.117 | r = -0.6 [-0.89, 0.02] |

| **Experiment 3** | | | | |
|---|---|---|---|---|
| Avoidance | Freezing | Two-way ANOVA | Sex: F(1, 23) = 0.41, p = 0.53<br>PA: F(1, 23) = 0.99, p = 0.33<br>S*PA: F( 1, 23) = 0.39, p = 0.54 | $\omega_p^2$ = -0.02 [0, 1]<br>$\omega_p^2$ < -0.01 [0, 1]<br>$\omega_p^2$ = -0.02 [0, 1] |
| Avoidance | Suppression of lever pressing | Non-parametric two-way ANOVA | Sex: F(1, 23) = 0.12, p = 0.73<br>PA: F(1, 23) = 0.49, p = 0.49<br>S*PA: F(1, 23) = 1.27, p = 0.27 | $\omega_p^2$ = -0.02 [0, 1]<br>$\omega_p^2$ = -0.04 [0, 1]<br>$\omega_p^2$ = -0.01 [0, 1] |
| Extinction | Freezing | Two-way ANOVA | Sex: F(1, 23) = 2.23, p = 0.15<br>PA: F(1, 23) = 2.69, p = 0.114<br>S*PA: F(1, 23) = 2.25, p = 0.147 | $\omega_p^2$ = 0.04 [0, 1]<br>$\omega_p^2$ = 0.06 [0, 1]<br>$\omega_p^2$ = 0.04 [0, 1] |
| Extinction | Suppression of lever pressing | Non-parametric two-way ANOVA | Sex: F(1, 23) = 1.79, p = 0.194<br>PA: F(1, 23) = 16.31, **p < 0.001**<br>S*PA: F(1, 23) = 0.78, p = 0.385 | $\omega_p^2$ = 0.01 [0, 1]<br>$\omega_p^2$ = 0.29 [0, 1]<br>$\omega_p^2$ = -0.02 [0, 1] |
